# The proteomics and phosphoproteomics landscape of melanoma under T cell attack

**DOI:** 10.1101/2025.09.12.675787

**Authors:** Giulia Franciosa, Agnete W. P. Jensen, Ana Martinez-Val, Ilaria Piga, Marco Donia, Jesper V. Olsen

## Abstract

Understanding how tumor cells interact with tumor-infiltrating lymphocytes (TILs) in the tumor microenvironment (TME) is crucial for identifying targetable immune checkpoints and predictive biomarkers for immunotherapy. While transcriptional responses have been characterized, protein-level changes remain largely unexplored.

Here, we used a system reproducing the interaction of TILs with cancer cells occurring in the TME, by co-culturing patient-derived cancer cells with matched autologous TILs at sub-lethal ratios. Using this system, we profiled the early response that cancer (melanoma) cells and TILs activate following autologous T cell attack. To distinguish melanoma from TIL proteomes, we applied stable isotope labeling by amino acids in cell culture (SILAC) combined with Orbitrap Astral-based data-independent acquisition (DIA) mass spectrometry, enabling cell type-specific profiling of protein and phosphorylation dynamics without FACS sorting. This approach also captured the global newly synthesized proteome of the mixed cultures.

Our analyses resolved interferon-γ-dependent proteome changes occurring in melanoma cells, identified the cytotoxic and regulatory T-cell molecule (CRTAM) as a selective marker of reactive cytotoxic T lymphocytes, and revealed tumor-intrinsic kinase activation signatures. Among these, multiple DNA damage response-associated kinases were activated during immune attack, suggesting potential therapeutic vulnerabilities.

Overall, this framework enables proteomic dissection of tumor–immune interactions and provides a resource for guiding biomarker discovery and therapeutic strategies to improve immunotherapy outcomes.

## Introduction

Enhancing a patient’s immune system through immune checkpoint blockade (ICB) can cure patients with metastatic melanoma and other malignancies. However, nearly half of patients with melanoma (1) and most with other solid tumors do not benefit from current ICB therapies, such as anti-CTLA-4, anti-PD1 and anti-PD-L1 (2). Treatment failure has been associated with a low number of immunogenic antigens, defective antigen presentation and/or the expression of alternative immune checkpoint molecules. Tumor-infiltrating lymphocytes (TILs) are specialized immune cells capable of targeting cancer cells, and their pre-existing activity within immune “hot” tumors is associated with a greater likelihood of ICB response. However, no consensus biomarkers currently exist to quantify intra-tumoral TIL activity and accurately predict ICB efficacy in individual patients. This highlights the urgent need for both improved therapeutic strategies and reliable predictive biomarkers for therapy response.

Previous efforts to identify resistance mechanisms or biomarkers in hot tumors have largely relied on genomics and transcriptomics (3–5). While informative, these approaches fail to capture early and dynamic signaling rewiring events mediated by protein post-translational modifications (PTMs). PTMs, particularly reversible and site-specific phosphorylation catalyzed by protein kinases, act as critical regulators of protein function and cellular signaling. In immunology, protein phosphorylation governs essential processes such as cytokine production, immune cell differentiation, and pathogen recognition, playing a central role in both innate and adaptive immunity. Mass spectrometry (MS)-based phosphoproteomics, the large-scale study of protein phosphorylation by LC-MS (6), enables the identification of ∼30,000 phosphorylation sites within just half-an-hour of data acquisition (7), providing a functional snapshot of cellular kinase and phosphatase activity.

We previously demonstrated that TILs isolated from melanoma patients remain reactive against autologous tumor cells in 2D co-cultures (8). However, applying phosphoproteomics to such co-culture systems presents a key challenge: cell lysis eliminates cell identity information, while manual separation is inefficient (3). Fluorescence-activated cell sorting (FACS) could be an alternative, but no published protocols exist for phosphoproteomics of sorted cells.

To address these limitations, we used stable isotope labeling by amino acids in cell culture (SILAC) (9) to discriminate tumor cells from TILs prior to MS analysis. Because newly synthesized proteins can be of either tumor or TILs origin during co-culture, we performed the co-culture in a medium containing a third stable isotope-labeled amino acid variant to capture cell type-specific signaling. This approach has previously been shown to provide biological insights into tumor-T cell networks (10). While SILAC has historically been combined with data-dependent acquisition (DDA), our approach uniquely integrates SILAC with data-independent acquisition (DIA) using the Orbitrap Astral mass spectrometer (11). Although SILAC or MS1-based multiplexing approaches and DIA have been successfully combined before (12–16), they have never been used as a method to distinguish different cell populations. This novel strategy allowed us to simultaneously analyze newly synthesized proteins (shared origin), as well as cell type-specific protein turnover and phosphorylation site changes with unprecedented depth.

## Methods

### Sample origin

All procedures were approved by the Scientific Ethics Committee for the Capital Region of Denmark. Tumor biopsies were obtained from four patients diagnosed with cutaneous metastatic melanoma, enrolled in the clinical trials at the National Center for Cancer Immune Therapy (CCIT-DK), Department of Oncology, Copenhagen University Hospital, Herlev, Denmark (H-20070020). Written informed consent was obtained from patients before any procedure according to the Declaration of Helsinki.

The four samples included in the study were derived from two anti-PD1 therapy-naïve and two patients with confirmed resistance to anti-PD1 therapy (**Supplementary Table 1**). Melanoma was chosen as a tumor model with known high TIL reactivity (17).

### Establishment of REP TILs and primary melanoma cell lines

From each biopsy, a matched pair of primary melanoma cells and autologous TILs were established *in vitro*, as described elsewhere (18, 19). Briefly, tumor specimens were obtained fresh and immediately transported to the laboratory in RPMI 1640 (Gibco, Thermo Fisher Scientific). The tumour masses were isolated from the surrounding tissues and tumours were sliced into multiple fragments (1-3 mm3 each) with a scalpel. Patient-derived melanoma cell lines were established from tumor fragments through short-term *in vitro* serial passages of adherent cells. After establishment, they were authenticated based on growth pattern, morphology and flow cytometry characterization analysis (described below). TILs were established *in vitro* by a two-step expansion process. First, they were “minimally expanded” (20) in high doses of IL2 (6,000 IU/mL) (Proleukin, Novartis) from tumor fragments. When a minimum of 50 x 10^6^ TILs were obtained (typically about 14-28 days after surgical resection), expansion was further achieved by a 14-days “rapid expansion” protocol (REP), in which TILs were unspecifically expanded with a 200-fold excess of allogeneic irradiated peripheral blood mononuclear cells (PBMCs) from healthy donors, high dose of IL2 and 30 ng/mL of anti-CD3 antibodies (clone OKT3, Miltenyi Biotec). The composition and functional reactivity of the REP TIL batches were subsequently analyzed via flow cytometry (described below).

### Cell culture, ligand stimulation and drug treatment

All cells were cultured at 37 °C in a humidified incubator with 5% CO2. Squamous Cell Carcinoma (SCC)-25 cells (male) were cultured in DMEM/H12 (Gibco, Thermo Fisher Scientific), supplemented with 10% fetal bovine serum (FBS, Gibco, Thermo Fisher Scientific), 100 U/ml penicillin and 100 μg/ml streptomycin (Pen-Strep, Gibco, Thermo Fisher Scientific). Primary melanoma cells were cultured in RPMI 1640 Medium with GlutaMAX (Gibco, Thermo Fisher Scientific), supplemented with 25 mM HEPES (Gibco, Thermo Fisher Scientific), 10% FBS and Pen-Strep. For short-term experiments, REP TILs were cultured in RPMI-1640 with GlutaMAX, supplemented with 25mM HEPES, 10% heat-inactivated human AB serum (HS, Sigma-Aldrich/Merck) and Pen-Strep. All cell lines were tested monthly for mycoplasma contamination by PCR.

SCC-25 were stimulated with 100 ng/ml of epidermal growth factor (EGF; Preprotech) for 8 minutes. Next, the cells were washed with PBS, trypsinized and centrifuged to obtain a cell pellet that was either lysed right away or incubated on ice for 3 hours with or without the following phosphatase inhibitors: 5 mM sodium orthovanadate, 1 mM sodium fluoride and 1 mM beta-glycerophosphate.

REP TILs were stimulated with: CD3/CD28 beads (Dynabeads™ Human T-Activator CD3/CD28, Gibco, Thermo Fisher Scientific) for 6 or 8 hours at 1 bead per 2 TILs ratio; CD3 (OKT3) antibody for 6 hours at a concentration of 30 ng/mL; 100 international units (IU) of recombinant interferon-γ (IFN-γ) for 6 hours; PMA/Ionomycin for 6 or 8 hours at a concentration of PMA 25 ng/mL and Ionomycin 0.5 µM.

Melanoma cells were stimulated with 100 IU of IFN-γ for 6 or 24 h.

### SILAC labeling of melanoma cells

Patient-derived melanoma cell lines were cultured in SILAC RPMI 1640 with GlutaMAX (Gibco, Thermo Fisher Scientific), 10% dialyzed fetal bovine serum (dFBS, Gibco, Thermo Fisher Scientific), Pen-Strep and 0.028 mg/mL of heavy L-Lysine-2HCl (^13^C ^15^N, Cambridge isotope laboratories) and 0.049 mg/mL of heavy L-Arginine-HCl (^13^C ^15^N, Cambridge isotope laboratories). After 10-14 days heavy isotope incorporation was assessed using MS. The resulting cell batches were cryopreserved for future use in co-culture experiments for proteomics analysis.

### Characterization of melanoma cell lines by flow cytometry

For flow cytometry analysis of melanoma cells, 5 x 10^5^ cells were collected and washed twice with Dulbecco’s phosphate-buffered saline (PBS, Sigma-Aldrich/Merck KGaA). Cells were then stained with Live/Dead™ Fixable Near-IR (APC-Cy7, Thermo Fisher Scientific) and subsequently for anti-MCSP (PE, clone EP-1, Miltenyi Biotec), anti-CD146 (BV421, clone P1H12, BioLegend) and anti-CD90 (FITC, clone 5R10, BioLegend) antibodies in PBS with 0.1% FBS at 4°C for 30 minutes, protected from light. After staining, cells were washed twice in PBS and analyzed on a NovoCyte Quanteon™ Flow Cytometer. Data was analyzed with NovoExpress software version 1.5.6 (Agilent).

### Flow cytometry analysis of activated TILs

Tumor-specific immune reactivity of REP TILs was assessed with 8-hour co-culture assays at 37°C with an effector:target ratio of 3:1. When assessing degranulation and cytokine production, the antibody anti-CD107a (BV421, Clone H4A3, BD Biosciences) was added prior to co-culture with brefeldin A (1:1,000, GolgiPlug, BD Biosciences) and monensin (1:1,000, GolgiStop, BD Biosciences). For CRTAM intracellular staining experiments, only monensin was added. REP TILs cultured alone or with PMA/Ionomycin served as negative and positive control, respectively. After the 8-hour co-culture, REP TILs were stained with Live/Dead™ Fixable Near-IR (APC-Cy7, Thermo Fisher Scientific) and the following antibodies for 30 minutes: anti-CD3 (BV786, clone SK7, BD Biosciences), anti-CD8 (Qdot 605, clone 3B5, Invitrogen; APC-R700 clone RPA-T8, BD Horizon), anti-CD4 (BV510, clone SK3, BD Biosciences), and anti-CRTAM (PE, clone Cr24.1 BioLegend). The cells were then washed, fixed and permeabilized overnight at 4°C using the FoxP3/Transcription Factor Staining Buffer Set (eBiosciences, Thermo Fisher Scientific). The following day the cells were stained with anti-CD137 (BV605, clone 4B4-1, BioLegend), anti-TNFα (APC, Clone MAb11, Invitrogen) and anti-IFNγ (PE-Cy7, Clone B27, BD Biosciences). Cells were analyzed on a NovoCyte Quanteon™ Flow Cytometer. Data was analyzed using NovoExpress software version 1.5.6 (Agilent). For CRTAM expression and co-expression experiments, cells were additionally extracellular stained with anti-CD137 (APC, clone 4B4-1; BioLegend) and CD25 (PE-Cy7, clone 2A3, BD Biosciences). Gates were defined using unstimulated TILs and fluorescence minus one (FMO) controls.

For flow cytometry experiments, wells from the same plate were considered technical replicates, while measurements performed on different days were considered biological replicates. All experiments were performed in three technical replicates.

### Real-time tumor killing analysis using xCELLigence

The cytotoxicity of REP TILs against tumor cells was assessed using a real-time cell analysis (RTCA) assay on the xCELLigence RTCA eSight™ system (Agilent), according to the manufacturer’s instructions. Briefly, tumor cells were seeded into a 96-well RTCA E-plate and incubated for 24 hours, until reaching a Cell Index value of 1. The day prior to the co-culture experiment, REP TILs were thawed and rested overnight in TILs media. On the day of the experiment, half of the tumor medium was replaced with either fresh medium (control) or medium containing autologous REP TILs. Cell Index (CI) measurements were recorded hourly for 72 hours after the addition of REP TILs, using the RTCA eSight Software Basic version 1.1.1 (Agilent) for data acquisition and analysis. CI was normalized to the time point where REP TILs were added, to obtain the Normalized Cell Index (NCI). The percentage of tumor cell killing was calculated by dividing the NCI of the co-culture by the tumor alone and multiplied by 100. For killing assays, wells from the same plate were considered technical replicates, while measurements performed on different days were considered biological replicates. For all experiments, each experimental condition was performed in at least 3 technical replicates.

### Co-culture of heavy-labeled melanoma cells with autologous REP TILs for proteomic and phosphoproteomic analysis

Prior to the co-culture experiment, REP TILs were thawed at rest overnight in TILs media supplemented with 6000 IU/mL of IL2. The following day, REP TILs were short-term-expanded in AIM-V supplemented with 10% HS and 6000 IU/mL of IL2 for 5 to 7 days. Two days before the co-culture, 1.5 x 10^6^ SILAC-heavy labeled tumor cells were seeded in a 10 cm petri dish in SILAC-heavy medium. After 24 hours, SILAC-heavy medium was replaced with serum-reduced SILAC-heavy medium (1% dFBS), while REP TILs were starved in AIM-V supplemented with 50 IU/mL of IL2. After 15 hours of starvation, 6 x 10^6^ REP TILs and ∼2 x 10^6^ tumor cells were processed for proteomic analysis and used as control samples. The rest of the seeded tumor cells were washed with 37 °C PBS, followed by incubation with serum-reduced SILAC medium-heavy media (SILAC RPMI 1640 with GlutaMAX™, 10% dFBS, Pen-Strep, 0.028mg/mL of L-Lysine-2HCl (4,4,5,5-D4, Cambridge isotope laboratories) and 0.049mg/mL L-Arginine-HCl (^13^C_6_, Cambridge isotope laboratories) with 2 x 10^6^ REP TILs. As a control, tumor cells and TILs were incubated separately. Samples were harvested before co-culture (0 hours), and after 2 and 6 hours. REP TILs were recovered from the supernatant, washed in PBS and centrifuged to obtain a cell pellet. Tumor cells were washed with PBS while still adhering on the plate. Match samples were reunited after lysis. For the proteomics experiment, each condition per patient was performed in six biological replicates (one replicate = one dish). Replicates were generated in close succession, within approximately one week. All samples downstream of cell lysis were processed and acquired together.

For the spectral library generation, this experiment was performed in one biological replicate without control samples and with unlabeled melanoma cells. For each patient, time points 2 and 6 hours were pulled.

### Sample preparation for proteomic and phosphoproteomic analysis

#### Cell lysis and protein extraction

Cells were lysed with boiling lysis buffer [5% sodium dodecyl sulfate (SDS), 5 mM tris(2-carboxyethyl)phosphine (TCEP), 10 mM chloroacetamide (CAA), 100 mM Tris HCl pH 8.5] and incubated for 10 min at 99 C while shaking. Samples were sonicated and protein concentration was determined using a BCA assay (Pierce).

#### Protein digestion

For each sample, 50-300 µg of protein was digested overnight with Lys C (FUJIFILM Wako Pure Chemical Corporation) and Trypsin (Sigma-Aldrich) at an enzyme:protein ratio of 1:500 and 1:250, respectively, using the KingFisher magnetic particle separation robot using the optimized protocol explained in detailed elsewhere (21, 22). The following day, samples were acidified with trifluoroacetic (TFA) to a final concentration of 1%. 0.5-1 µg of peptides were loaded on Evotips (Evosep Biosystems) for proteome analysis. The rest of the peptides were further processed by desalting with solid-phase extraction using C18 Sep-Paks (Waters Corporation), followed by speedvac until acetonitrile (ACN) evaporation. The final peptide concentration was estimated by measuring absorbance at 280 nm on a NanoDrop 2000C (Thermo Scientific), before proceeding with Ti-IMAC phosphopeptide enrichment.

#### High-pH reversed-phase peptide fractionation

To generate a spectral library for patient 905, 200 μg of unlabeled peptides were resuspended in 20 μl of 25 mM ABC and fractionated using a reversed-phase Acquity CSH C18 1.7 μm x 1 mm × 150 mm column (Waters) coupled to the UltiMate 3000 high-performance liquid chromatography (HPLC) system (Thermo Fisher Scientific) by using the Chromeleon software (Thermo Fisher Scientific). The instrument operated at 30 μl/minute with column oven temperature set to 40 °C. Buffer A (5 mM ABC) and buffer B (100% ACN) were used. Peptides were eluted in 12 fractions with concatenation via multi-step gradient as follows: 0-62.5 min 8-28% B; 62.5-67 min 28-60% B; 67-70 min 60-70% B; 70-77 min 70% B; 77-78 min 8% B; 78-87 min 8% B. Fractions were acidified with formic acid (FA; 40 μl of 10% FA) to a final concentration of 1% and phosphorylated peptides were enriched by Ti-IMAC enrichment.

#### Enrichment of phosphorylated peptides

Ti-IMAC phosphopeptide enrichment was carried out on a KingFisher Flex robot (Thermo Fisher Scientific) in 96-well format, as previously described (21–23). 18 μg of peptide was used as input for enrichment, with 5 μl of magnetic Zr-IMAC HP beads (ReSyn Biosciences). Eluted phosphopeptides were acidified with 40 μl of 10% TFA and loaded on Evotips (Evosep Biosystems) for MS analysis.

### LC-MS/MS analysis

LC-MS/MS analysis was performed on an Orbitrap Astral mass spectrometer (Thermo Fisher Scientific) (11) coupled to a Vanquish Neo UHPLC (Thermo Fisher Scientific) in the trap-and-elute mode or an Evosep One LC system (Evosep Biosystems) (24), and interfaced online using an EASY-Spray source. The analytical column type was chosen according to the experiment (**Supplementary Table 2**).

The Orbitrap Astral mass spectrometer was operated in positive ion mode with data independent acquisition (DIA). The spray voltage was set at static, 1.8 or 2 kV, the heated capillary temperature at 275 or 280 °C and funnel RF frequency at 40. The full-MS resolution was set at 120K, 180K or 240K with a full scan range of m/z 380–980 for proteome and m/z 480-1080 for phosphoproteome. The full-MS automatic gain control (AGC) was set to 500% and the maximum injection time was set at 3, 5, 10 or 30 ms. The isolated peptide precursor ions were fragmented using HCD with 25% Normalized Collision Energy (NCE). The fragment scan range was set at 150-2000 m/z. DIA-MS/MS fragment ion scans were recorded with a 2, 4 or 6 Th quadrupole isolation window and a maximum injection time of 2.5, 3, 6 or 8 ms. The MS method details for each experiment are described in **Supplementary Table 2**.

To increase the depth of the project-specific phosphoproteomics spectral library, the unlabeled co-culture samples from patient 905 were subjected to high-pH fractionation, phosphopeptide enrichment and MS analysis, as described above. Additionally, samples from all patients were phosphoenriched without prior high-pH fractionation and analysed by gas-phase fractionation during MS analysis. Details about the MS method are found in **Supplementary Table 2**.

### Raw mass spectrometry data processing

Raw MS data were analyzed using DIA-NN (25), version 2.0. The Human Uniprot fasta file was downloaded in October 2024 and contained 20,428 entries. Two spectral libraries (one for proteome and one for phosphoproteome) were generated *in-silico* by enabling “FASTA digest for library-free search / library generation” and “Deep learning-based spectra, RTs and IMs prediction”. No raw data was supplied in this step. For the proteome library, the settings were the following: maximum number of missed cleavages set to 1, maximum number of variable modifications set to 0, N-terminal methionine excision enabled, cysteine carbamidomethylation enabled as a fixed modification, minimum peptide length set to 7, maximum peptide length set to 30, minimum precursor charge set to 2, maximum precursor charge set to 4, minimum precursor m/z set to 380, maximum precursor m/z set to 980, minimum fragment m/z set to 150, max fragment m/z set to 2000, contaminants enabled. For the phosphoproteome library, the settings were the following: maximum number of missed cleavages set to 2, maximum number of variable modifications set to 3, N-terminal methionine excision enabled, cysteine carbamidomethylation enabled as a fixed modification, phosphorylation on STY enabled as variable modification, minimum peptide length set to 7, maximum peptide length set to 30, minimum precursor charge set to 2, maximum precursor charge set to 4, minimum precursor m/z set to 480, maximum precursor m/z set to 1080, minimum fragment m/z set to 150, max fragment m/z set to 2000, contaminants enabled.

The “in-silico” spectral libraries were supplied to search raw MS data with the same settings used for the libraries, except for “FASTA digest for library-free search / library generation” and “Deep learning-based spectra, RTs and IMs prediction” that were unchecked. Mass accuracy and scan window were fixed (values depend on the experiment: see **Supplementary Table 2**). When searching SILAC raw files, the following text was added to “Additional options”:

--fixed-mod SILAC,0.0,KR,label

--lib-fixed-mod SILAC

--channels SILAC,L,KR,0:0; SILAC,M,KR,4.025107:6.020129;SILAC,H,KR,8.014199:10.008269

--original-mods

--channel-spec-norm

For the SILAC proteome search, “generate spectral library” was enabled and MBR was not enabled. The obtained empirical spectral library was supplied to search raw MS data separately for each time point, gradient length and MS method, using the same settings discussed above, except for “Additional options”, where --lib-fixed-mod SILAC was not added.

For the search of the project-specific phospho-enriched fractionated spectral library, “generate spectral library” was enabled, “Unrelated runs” was enabled and MBR was not enabled. The obtained spectral library was supplied to search SILAC phosphoproteomic raw MS data separately for each time point.

For the SCC-25 phosphoproteome and the IFN-γ proteome searches, MBR was enabled.

### Bioinformatic analysis

All bioinformatic analysis was performed using R version 4.4.1 with R studio version 2024.12.1.

#### Proteomics data filtering and processing

For all SILAC datasets, the main report in .parquet format was used and processed with the R package “arrow”. For the IFN-γ proteome dataset, “pg_matrix.tsv” was used. For the SCC-25 phosphoproteome dataset, “phosphosites_90.tsv” was used.

For all datasets, contaminants and entries without gene names were removed, and MS intensities were log2 transformed.

For all SILAC datasets, entries without a channel were removed. Data were filtered at 1% FDR, using global q-values for protein groups (“Lib.PG.Q.Value” <= 0.01) and both global and run-specific q-values for precursors (“Q.Value” and “Lib.Q.Value” <= 0.01, respectively). An additional 5% run-specific protein-level FDR filter (“PG.Q.Value” <= 0.05) was applied too.

For all SILAC proteome datasets, only protein entries with a “PG.MaxLFQ.Quality” >= 0.7 were kept. The column “PG.MaxLFQ” was used for downstream quantitative analyses.

For the SILAC phosphoproteome dataset, data were filtered at 1% FDR also using global and run-specific peptidoform q-values (“Peptidoform.Q.Value” and “Lib.Peptidoform.Q.Value” <= 0.01, respectively), and channel-level q-value (“Channel.Q.Value” <= 0.01). Only precursors with a “Quantity.Quality” >= 0.5 that were phosphorylated on a serine, threonine or tyrosine were kept. The column “Precursor.Normalised” was used for downstream quantitative analyses. Precursor-level quantifications were aggregated to the phosphosite level based on the “Protein.Sites” column, considering only phosphorylation events and excluding other modifications. Phosphosites detected on precursors with different phosphorylation multiplicities (i.e., singly, doubly, or triply phosphorylated) were not merged and were labeled accordingly: M1 for singly, M2 for doubly and M3 for triply phosphorylated precursors. Run-specific site localization probabilities were extracted from the “Site.Occupancy.Probabilities” column. Prior to aggregation, any phosphosite with a localization probability below 0.75 in a given run was excluded from the analysis. The aggregation was performed using the “top-1 method”: for each site, the precursor entry with the highest summed intensity across runs was retained. The sequence window was extracted from the protein sequence, including 5 amino acids before and after the phosphorylated residue. The protein sequence was retrieved with the R package “Protti”, using the function “fetch_uniprot”.

Proteins belonging to the EGFR signaling pathway were retrieved using the R package “KEGGREST” as part of the KEGG pathway “ErbB signaling pathway” (hsa04012).

Labeling efficiency for the heavy channel was calculated at the time point zero-hours for the tumor-alone samples as the number of protein groups identified in the heavy channel divided by the total number of protein groups identified across all channels.

Heavy channel false discovery rate (FDR) was calculated at the time point zero-hours for the TILs-alone samples as the number of protein groups identified in the heavy channel divided by the total number of protein groups identified across all channels. Medium FDR was calculated at the time point 0-hours separately for tumor- and TILs-alone samples as the number of protein groups identified in the medium channel divided by the total number of protein groups identified across all channels.

Protein group coefficient of variation (CV) was calculated only for the co-culture samples by dividing the standard deviation (SD) by the mean of raw MS intensity for entries with at least 3 out of 4 valid values per channel. Pearson correlation was calculated using pairwise complete observations.

Principal component analysis (PCA) was performed with entries identified across the entire dataset.

#### Differential expression analysis (DEA)

Data was further filtered before DEA. For the SCC-25 data, only entries with at least 3 valid values in at least one experimental group (lL_Ctrl, IL_EGF, PBS_Ctrl, PBS_EGF, PI_Ctrl, PI_EGF) were included. For the SILAC proteome and phosphoproteome datasets, DEA was performed separately on each channel. For both datasets and the IFN-γ proteome dataset, only entries with at least 2 valid values in at least 3 patients and 3 valid values per experimental group overall (control and co-culture/IFN-γ) were included. For the SILAC phosphoproteome dataset, DEA was also performed on each individual patient and channel. Only entries with at least 2 valid values per experimental group (control and co-culture) were included.

DEA was performed using the limma package (26, 27). We accounted for repeated measures by using the “duplicateCorrelation” function to estimate the correlation between samples from the same patient. This correlation was incorporated into the model via the “block” argument in the “lmFit” function (28).For the SILAC phosphoproteome dataset, DEA was also performed per patient and channel, and no blocks were set. For the SCC-25 phosphoproteome dataset, no blocks were set. Empirical Bayes moderation was applied using “eBayes” with “trend = TRUE”, which allows the prior variance estimate to depend on the average intensity of each protein (or phosphosite) (29). P values were corrected using the Benjamini Hochberg (BH) procedure. An entry was considered regulated if the adjusted p value was equal or smaller than 0.01.

A fold-change cutoff was also applied to define significant regulation. To define a cutoff in the SCC-25 phosphoproteome, SD was calculated separately for each contrast (IL, PBS, PI), and for up- and down-regulated phosphosites. This value multiplied by 2 ranged from 0.9 to 1. Therefore, a phosphosite was considered regulated if the log2 fold-change was equal or greater than 1 for up-regulation and -1 for down-regulation. In the IFN-γ proteome, the SD was calculated separately for each time point (6 and 24 hours), and for up- and down-regulated proteins. This value multiplied by 2 was used as a fold-change cutoff (6 hours: up-regulation = 0.24; down-regulation = -0.26; 24 hours: up-regulation = 0.75; down-regulation = -0.37). As in the SILAC proteome dataset the fold-change distribution was tailing towards the co-culture side, we employed the median absolute deviation (MAD) instead of the SD. MAD was calculated separately for each channel, time point, and for up- and down-regulated proteins. The mean of these 6 values multiplied by 5 was used as a fold-change cutoff (2 hours: up-regulation = 0.3; down-regulation = -0.3; 6 hours: up-regulation = 0.35; down-regulation = -0.35). In the SILAC phosphoproteome dataset, MAD was calculated separately for each channel, and for up- and down-regulated proteins. The mean of these 6 values multiplied by 5 was used as a fold-change cutoff (up = 0.65, down = -0.65). When performing DEA on the SILAC phosphoproteome dataset per patient and channel, no fold-change cutoff was applied.

#### Over-representation analysis (ORA)

ORA was performed on proteome data using the R package “clusterProfiler” (30). All the proteins identified in the medium-heavy channel were used as background. For gene ontology biological process ORA, Uniprot ids were converted to Entrez ids, and only 299 out of 301 had a match.

#### Single-sample gene set enrichment analysis (ssGSEA)

ssGSEA (31) was performed on GenePattern using the standard settings.

#### Motif enrichment and visualization analysis

Motif visualization analysis was performed using the pLogo generation tool (32), publicly available at http://plogo.uconn.edu/. As foreground, we used the sequence window of sites significantly up-regulated upon attack; as background, we used the whole identified phosphoproteome in the heavy channel (after the initial filtering steps). “No fixed position” was selected.

#### Kinase-substrate and pathway enrichment analysis

Kinase-substrate enrichment analysis was performed with RoKAi v2.3.0 (33) and with PTM-Signature Enrichment Analysis (PTM-SEA) (34), the latter as implemented in PTMNavigator (35). To select kinases from the PTM-SEA output, only signature IDs containing the string “KINASE” were kept. Pathway enrichment analysis for phosphoproteomics data was performed with redundant single-sample Gene Set Enrichment Analysis (ssGSEA) 2.0 or (31) with PTM-Signature Enrichment Analysis (PTM-SEA), both implemented in PTMNavigator. We ranked sites using a signed -log10(p-value) metric derived from the Limma output. Specifically, for each phosphosite, the -log10 of the p-value was used if the corresponding log2 fold change was positive, while the log10 of the p-value (i.e., a negative value) was used if the fold change was negative. For the SILAC phosphoproteome data, phosphosites with the same residue and position but different multiplicity were collapsed to one value by keeping the maximum absolute value.

### STRING network analysis

The functional protein network displayed in Figure 3C was generated in Cytoscape using STRING (36) as implemented in the STRING app (37). The network was visualized using the Omics Visualizer app (38).

### CRISPR-KO screens

Data were downloaded from the original publications (39, 40). In the Kearney et al. dataset, genes were considered positively enriched if their positive p value was <= 0.01, while they were considered negatively enriched if their negative p value was <= 0.01. In the Zhang et al. dataset, genes were considered positively enriched if their positive fdr was <= 0.05, while they were considered negatively enriched if their negative fdr was <= 0.05. Enriched genes in either dataset were considered enriched and used for the analysis shown in Figure S11.

### Kaplan-Meier (KM) survival analysis

The KM curves shown in Figure 5E and S17C were made with the Kaplan Meier plotter tool at https://kmplot.com/analysis/ using the immunotherapy tab. Tumor type was restricted to melanoma, while all the other settings were left unchanged. The KM curve shown in Figure S16B was made in TIDE (see paragraph below).

### Single-cell RNA sequencing (ssRNA-seq)

ssRNA-seq data (41) were downloaded from the <u>Single Cell Portal</u>. Data was split in quartiles based on CRTAM gene expression. 0s were considered a separate group. Difference in the expression of selected genes between CRTAM quartiles was assessed by pairwise Wilcoxon Rank Sum. p values were adjusted with the Benjamini-Hochberg procedure.

### TIDE (Tumor Immune Dysfunction and Exclusion) analysis

Genes of interest were queries on the “Query Gene” tab on the <u>TIDE website</u>, and the data were downloaded from the “Expression” tab. Cancer types with less than 3 cohorts were excluded. Only overall survival (OS) data were used.

### Data visualization

Plots were performed using the R package “ggplot2” version 3.5.1 or the GraphPad Prism software version 10. Heatmaps were generated through the R package “pheatmap” version 1.0.12 or the R package “ggplot2” version 3.5.1. Euler diagrams were made through the R package “eulerr”.

## Results

### *In vitro* recognition of patient-derived melanoma cells by autologous T cell attack

To study the early signaling events during T cell-mediated attack of melanoma cells, we used a patient-derived 2D co-culture system in which melanoma cell lines established from tumor biopsies are targeted by autologous TILs isolated from the same biopsies (8). The tumor:TIL pairs were selected based on high anti-tumor reactivity and efficient tumor cell killing in preliminary assays. Flow cytometric assessment of anti-tumor reactivity using an effector-to-target (E:T) ratio of 3:1 confirmed a robust TIL activation across all selected pairs, demonstrated by the upregulation of the costimulatory molecule CD137 (42), the degranulation marker CD107a (43), and the increased secretion of IFNγ and TNFα (**Figure 1A**, **S1** and **S2A**). Since we aimed to capture early signaling events during tumor cell killing while maintaining comparable cytotoxic capacity across all four tumor:TIL pairs, we performed E:T ratio titrations (**Figure S2B**) to identify conditions that resulted in equivalent tumor cell killing without inducing excessive cell death at early time points, which will represent a more natural interaction of TILs and cancer cells *in vivo*, and would not compromise downstream MS analyses. Based on these titrations, an E:T ratio of 1:1 for all patients was selected for subsequent MS experiments (**Figure 1B**). Prior to performing any experiment, melanoma cell lines were analyzed by flow cytometry to confirm minimal fibroblast contamination (**Figure S2C**). Additionally, TILs were characterised for the relative abundance of CD8^+^ and CD4^+^ T cells, highlighting that three of the four TIL products (P905, P915, P924) were predominantly CD8^+^ T cells, whereas one (P605) contained approximately equal proportions of CD8^+^ and CD4^+^ T cells (**Figure S2D**).

**Figure 1.**
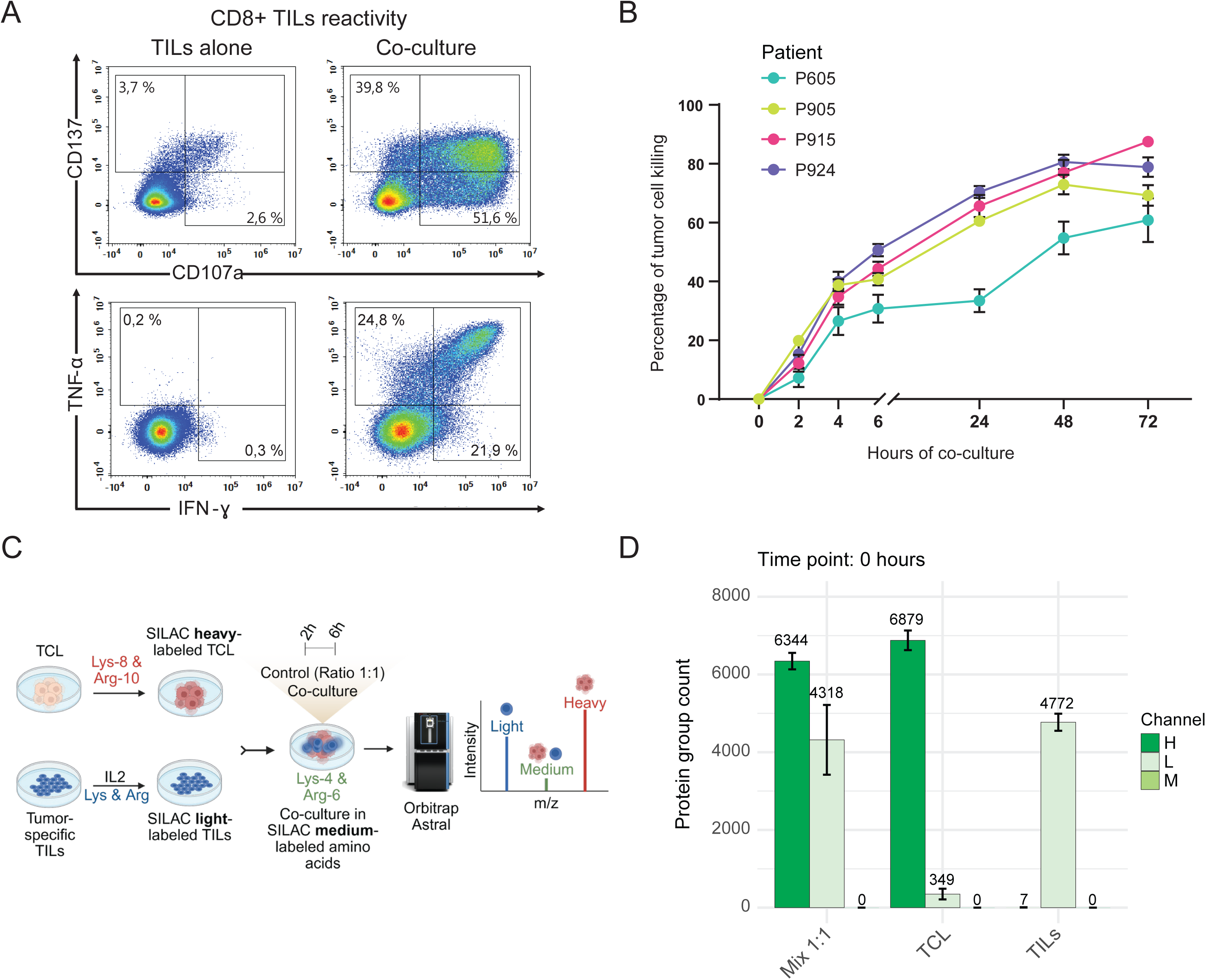
SILAC-DIA proteomic analysis of patient-derived melanoma cells under attack by autologous TILs. **A**. Representative density plots of flow cytometric analysis of surface expression of CD137 and CD107a, and intracellular expression of TNF-α and IFN-γ in activated CD8^+^ TILs from patient 905. TILs were co-cultured for 8 hours with autologous melanoma cells at effector:target ratio 3:1. **B**. Autologous TIL-mediated killing of melanoma cells from four different patients by xCELLigence real-time cell analysis at an effector:target ratio of 1:1. The percentage of tumor cell killing was calculated as the Normalized Cell Index (NCI) of the tumor in coculture divided by the NCI of tumor alone x 100. Data are presented as mean ± SD (n=3-4 technical replicates). **C**. Schematic drawing of the proteomics pilot experiment performed in patient 905. Created with Biorender. **D**. Protein group identifications per channel and cell type at time point 0-hours (before co-culture). Data are presented as mean ± SD (n=4 biological replicates). Samples were analyzed on an 180 sample-per-day gradient.

### Loss of phosphorylation dynamics due to prolonged sample processing

To develop an optimal method for analyzing phosphorylation site dynamics in melanoma cells under T cell attack using MS-based phosphoproteomics, we first evaluated whether fluorescence-activated cell sorting (FACS) could be used to physically separate tumor cells and TILs while preserving their *in vivo* phosphorylation state prior to sample preparation and MS analysis. As a model system, we used SCC-25 squamous cell carcinoma cells stimulated with recombinant epidermal growth factor (EGF) for 8 minutes. Cells were lysed immediately after stimulation using a hot lysis buffer to preserve phosphorylation by inactivating proteases and phosphatases. To simulate FACS-related sample processing, cells were incubated on ice for three hours prior to lysis, with or without the addition of phosphatase inhibitors (**Figure S3A**). This experiment showed that while the total number of identified phosphorylation sites was only slightly reduced by the prolonged incubation, this effect was negligible in the presence of phosphatase inhibitors (**Figure S3B**). However, most of the significantly EGF-regulated phosphosites lost their regulation following incubation on ice, and this effect was surprisingly even more pronounced when phosphatase inhibitors were added (**Figure S3C**). When focusing on the EGFR signaling pathway, we found that the dynamic regulation of key phosphosites involved in the pathway was severely compromised by prolonged incubation (**Figures S3D**). Phosphatase inhibitors did not rescue this loss of regulation and, in some cases, even led to inverse regulation. When performing PTM-Signature Enrichment Analysis (PTM-SEA), prolonged sample handling markedly reduced the number of significantly regulated pathways (**Figure S3E**), with EGFR signaling entirely abrogated in PBS-treated samples and only partially retained in the presence of phosphatase inhibitors (**Figure S3F**).

These findings suggest that prolonged processing steps such as FACS sorting interfere with the detection of dynamic phosphorylation changes. Therefore, they are not compatible with phosphoproteomics methods aimed at capturing signaling responses.

### SILAC-DIA for cell type-specific phosphoproteomic signatures

Because FACS could not be used, cell type-specific information had to be obtained without physically separating tumor cells and TILs prior to lysis. To achieve this, we employed SILAC labeling of tumor cells using “heavy”-labeled amino acids (K8, R10), while TILs were left unlabeled (K0, R0). The two cell types were co-cultured for 2 and 6 hours, respectively, in media containing “medium-heavy”-labeled amino acids (K4, R6). This approach ensures that newly synthesized proteins, which cannot be assigned to a specific cell type, may be excluded from downstream analysis. As a control, tumor cells and TILs were incubated separately and mixed immediately before lysis (**Figure 1C**). A 1:1 tumor-to-TIL ratio was used, based on prior optimization (**Figure 1A-B** and **S2B**).

To test the quantitative performance of the SILAC strategy, we selected one patient pair (P905) to perform a pilot experiment in four biological replicates. Full proteome samples were analyzed on the Orbitrap Astral mass spectrometer using narrow-window DIA (nDIA) (11) and short LC gradients, and the resulting MS data was processed with DIA-NN 2.0 (25) using the plexDIA module (15) coupled with the QuantUMS (Quantification using an Uncertainty Minimising Solution) quantification method (44). Tumor cells and TILs were also collected and analyzed separately at time point zero-hours, before adding the medium-heavy amino acids. TIL-only samples yielded almost exclusively light-labeled peptide identifications, with a calculated mean false discovery rate (FDR) in the heavy channel of 0.1% (**Figure 1D**). Slightly more light-labeled peptides were detected in tumor samples, likely due to a not full incorporation of heavy amino acids. Importantly, no medium-heavy-labeled peptides were detected in either cell type at zero-hours prior to co-culture, yielding a calculated mean FDR in the medium-heavy channel of 0%. This confirmed the effectiveness of the plexDIA module for our experimental setup.

Using the SILAC-based co-culture approach, we next asked whether tumor cells and TILs could simply be separated after co-culture to obtain clean, cell type-specific proteomes. If this were feasible, SILAC labeling would not be required. However, manual separation at both 2 and 6 hours resulted in substantial cross-contamination, with “light” peptides detected in tumor cell fractions and vice versa, already evident at the 2-hour time point (**Figure S4A-B**). These results confirmed previous observations that post-co-culture separation of interacting cell populations is insufficient, thereby justifying the need for the SILAC-based strategy (3).

Quantification in DIA-NN 2.0 leverages both MS1 and MS2 data to enable precursor quantification (44). To evaluate whether the MS2-oriented nDIA method (Method 1) (11) was the most optimal for multiplexed DIA in this context, we compared it to an MS1-oriented nDIA method (Method 2). Given that TILs are smaller and contribute less protein than tumor cells in the mixed tumor-TILs sample, we also tested whether a wider DIA window (Method 3) would improve ion accumulation and detection depth for TIL-derived peptides. Among the three, Method 1 outperformed the others, yielding the highest number of identifications (**Figure S4C**) and the lowest coefficients of variation (CV) (**Figure S4D**), serving as a proxy for quantification precision.

### Proteomic and phosphoproteomic analysis of melanoma-TIL pairs

To comprehensively investigate the signaling dynamics underlying the interaction between patient-derived melanoma cells and TILs, we performed a large co-culture experiment using cells from all four patients (**Figure 2A-B**) for both proteomic and phosphoproteomics analysis. Each condition was assessed in six biological replicates at two time points (2 and 6 hours). In addition, tumor cells and TILs were collected and analyzed separately at time point zero-hours for proteome analysis to (i) verify the success of stable isotope labeling (tumor: heavy; TILs: light) and (ii) increase the depth of the empirical DIA library (see methods). The labeling efficiency was higher than 90% in all cell lines: 97% for P605, 94% for P905, 97% for P915 and P924. Light channel FDR ranged from 0 to 0.4%, while medium-heavy channel FDR was 0% across all patients (**Figure S5A**).

**Figure 2.**
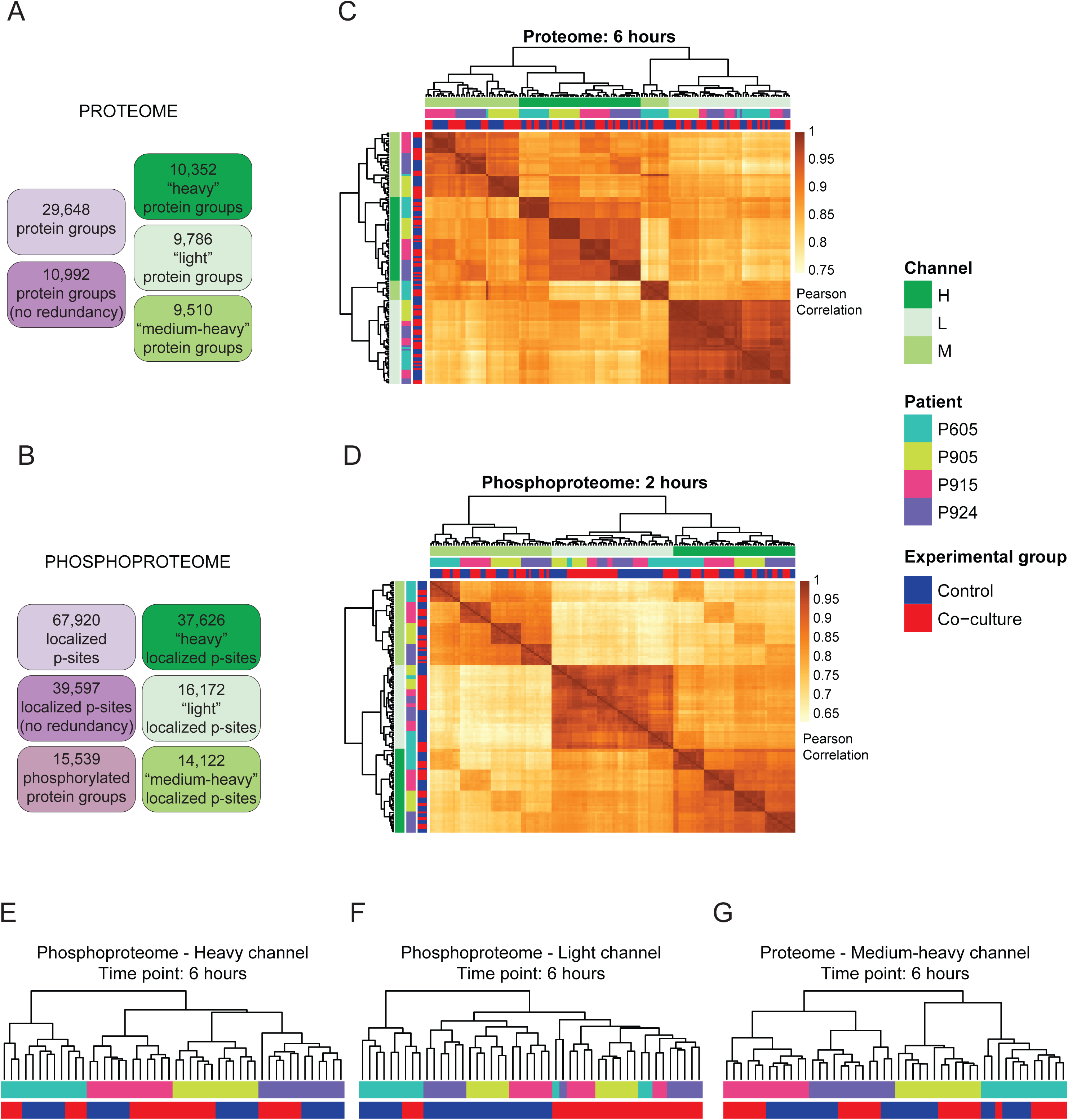
Proteomic and phosphoproteomic analysis of four melanoma-TIL pairs. **A-B**. Number of identifications in the proteomics and phosphoproteomics experiment, respectively. **C-D**. Pearson correlation heatmaps of Log2 MS intensities, including all 96 samples analyzed in the proteomics (6 hours) and phosphoproteomics (2 hours) experiment, respectively. **E-G**. Dendrogram representation of hierarchical clustering with euclidean distance of Log2 MS intensities for the phosphoproteome of the heavy (E) and light channels (F), and the proteome of the medium-heavy channel at 6-h time point (G).

Across all samples, we confidently identified approximately 30,000 protein groups in the proteome data. Of these, 10,352 were detected in the heavy channel, 9,786 in the light channel and 9,510 in the medium-heavy channel. Approximately 11,000 protein groups were identified regardless of channel assignment (**Figure 2A**); 10,175 were shared between at least two channels and 8,481 shared across all three (**Figure S5B**).

Due to the inherent size differences between melanoma and T cells, the light channel (TILs) exhibited the highest proportion of missing values. On average, 8,618 and 8,048 protein groups were identified per run in the heavy channel at 2 and 6 hours, respectively, while the light channel yielded an average per run of 4,151 and 3,715 identifications at the same time points. In the medium-heavy channel, which captures newly synthesized proteins, the number of identifications was initially low (2,645 per run at 2 hours), consistent with the detection of fast-turnover proteins only. However, by 6 hours, the average number of identifications rose to 6,211 per run on average, indicating that a substantial portion of the proteome had undergone synthesis and was incorporated into the medium-heavy label by this time (**Figure S5C**).

Given the lowest rate of protein turnover, we selected the 2-hour time point for phosphoproteomic analysis to focus on cell type-specific signaling in pre-existing (“old”) proteins, minimizing interference from the few newly synthesized proteins labeled in the medium-heavy channel. To increase the speed of the search and the phosphoproteome depth, we generated a project specific spectral library by performing the same co-culture experiment with unlabeled melanoma cells. Samples were processed through low input phosphoproteomics (23). Across all samples, we confidently identified approximately 70,000 localized phosphorylation sites (localization probability above 0.75) on 15,539 protein groups. Of these phosphosites, 37,626 were detected in the heavy channel, 16,172 in the light channel and 14,122 in the medium-heavy channel. Approximately 40,000 phosphorylation sites were identified regardless of channel assignment (**Figure 2B**); 19,290 were shared between at least two channels and 9,033 shared across all three (**Figure S5D**). On average, 16,157 and 8,048 phosphorylation sites were identified per run in the heavy channel, while the light channel yielded an average of 5,037 identifications (**Figure S5E**).

We generated Pearson correlation heatmaps for the proteome data at 6 hours (**Figure 2C**) and for the phosphoproteome data at 2 hours (**Figure 2D**). In all heatmaps, the main clustering was driven by the SILAC channel, confirming that our multiplexing approach effectively distinguishes both the cellular origin (tumor cells vs TILs) and the fraction of newly synthesized proteins. At 2 hours, the medium-heavy channel clustered separately from the heavy and light channels, whereas at 6 hours, it clustered together with the heavy channel, suggesting that the majority of newly synthesized proteins at later time points originate from tumor cells (which are larger and thus contain more protein). The only exception were samples from patient 605 whose medium-heavy channel was clustered with the light channel suggesting T cells are producing more proteins in this system, probably because of the highest prevalence of CD4 positive TILs (**Figure S2D**). Within the heavy and medium-heavy channels, samples were always clustered by patient, with three patients (P905, P915, P924) clustering separately from patient P605. Interestingly, TILs did not show a marked patient-specific clustering. We observed a certain level of clustering by condition (control vs. co-culture) in the heavy and light channel for the phosphoproteome at 2 hours and in the medium-heavy channel for the proteome at 6 hours. This separation improved when we increased the number of quantified sites included and used Euclidean distance, which allows for missing values, as the clustering metric (**Figure 2E-G**).

### Proteome changes upon T cell attack

On the proteome data, we performed differential expression analysis to identify proteins significantly regulated upon T cell attack in the same direction in all models. Proteins showing regulation in the heavy or light channels reflect cell type-specific changes and are primarily interpreted as differentially degraded upon co-culture, though a minor contribution from newly synthesized proteins incorporating recycled amino acids cannot be excluded. In contrast, proteins regulated in the medium-heavy channel represent changes in protein synthesis upon co-culture and may originate from either tumor cells or TILs. Most of the protein abundance changes were observed in the medium-heavy channel at 6 hours, with 263 up-regulated proteins and 80 down-regulated (**Figure S6A**), while few proteins were regulated at the 2-hour time point (**Figure S6B**) and in the heavy and light channels at the 6-hour time point (**Figure S6A**).

Proteins showing up-regulation in the medium-heavy channel showed a high degree of similarity across the four different models (**Figure 3A**). Since many immune-related proteins may exhibit a high number of missing values in the control condition of the medium-heavy channel, we systematically searched for other proteins with such a pattern, and identified 38 proteins preferentially expressed upon co-culture (**Figure 3B**) and 78 preferentially not expressed (data not shown). Among proteins up-regulated or exclusively induced upon T-cell attack, we identified the adhesion molecules ICAM1 and VCAM1 that may facilitate stable immune synapse formation, enabling cytotoxic T lymphocyte (CTL)-mediated killing, as supported by the presence of PRF1 and GZMB, key components of the CTL arsenal. We also found several known interferon-γ (IFN-γ) targets, for instance STAT1 and IRF1. At the same time, immune checkpoint proteins CD274 (PD-L1) and IDO1 emerged, suggesting attempts by tumor cells to resist immune attack.

**Figure 3.**
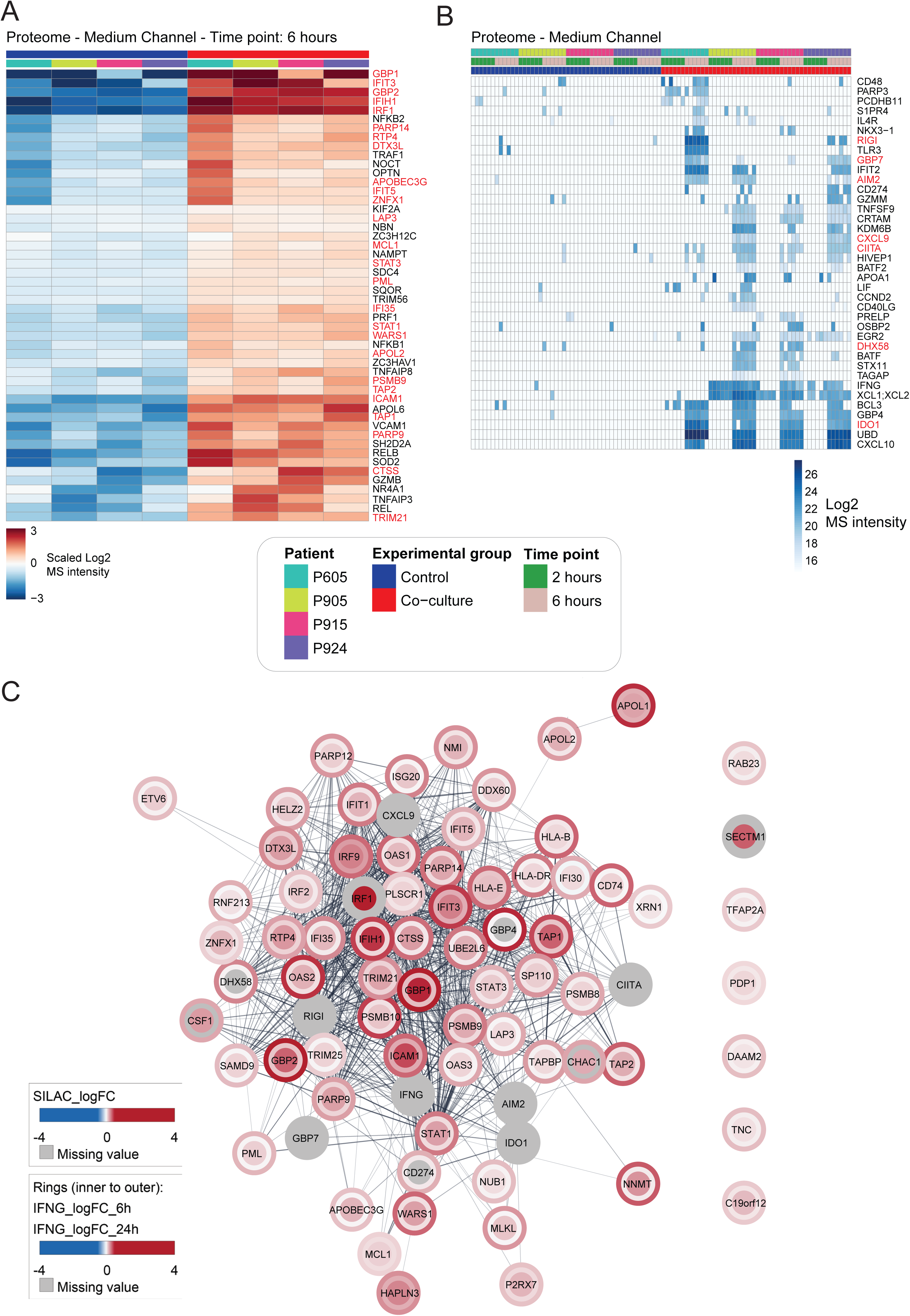
Proteome changes upon T cell attack in the medium-heavy channel. **A**. Heatmap of scaled log2 MS intensities of the 50 up-regulated proteins with the lowest adjusted p value in the medium-heavy channel at 6 hours. Missing values were imputed with the minimum value of the filtered matrix before scaling. Scaling was performed within each protein group and patient. The six replicates were averaged after scaling. Rows were clustered with euclidean distance, while the column order was fixed. Proteins labeled in red are induced by IFN-γ (therefore also displayed in C). **B**. Heatmap of Log2 MS intensities of the 38 proteins preferentially expressed upon co-culture, defined as having at most 2 out of 48 values in the control condition and at least 7 out of 48 in the co-culture condition. Missing values were imputed with a low value. Rows were clustered with euclidean distance, while the column order was fixed. Proteins labeled in red are induced by IFN-γ (therefore also displayed in C). **C**. Functional STRING protein network of the proteins up-regulated both by T cell attack in the medium-heavy channel at 6 hours and by IFN-γ stimulation of melanoma cells either at 6 or 24 hours. The selection criteria for which proteins are included in the network are explained in Figure S9A.

Activation of both canonical (NFKB1) and non-canonical (NFKB2 and RELB) NF-κB pathways, along with regulators like TNFAIP3, reflected sustained inflammatory signaling. Innate immune sensors including RIG-I (DDX58), DHX58, TLR3 and AIM2 were also upregulated, pointing to nucleic acid sensing triggered by immune-mediated stress. Transcription factors such as BATF, EGR2 and HIVEP1 indicated extensive transcriptional remodeling, while co-stimulatory and checkpoint-related proteins like TNFSF9 (4-1BBL), CD40LG and CD48 highlighted dynamic immune-tumor crosstalk. Finally, proteins such as NAMPT, SOD2 and OPTN suggested metabolic and oxidative adaptations that help tumor cells cope with the inflammatory microenvironment. Over-representation analysis (ORA) confirmed that these proteins were enriched only in immune related GO terms and pathways (**Figure S6C-D**).

### Deconvolution of the interferon-γ-dependent proteome changes upon T cell attack

We have previously shown that T cell attack induces broader transcriptomic changes in tumors compared to IFN–γ in melanoma (3). To confirm this on the proteome level, we stimulated three different patient-derived tumor cell lines (P905, P915, P924) with recombinant IFN-γ for 6 and 24 hours for MS-based proteomic analysis. Each condition was performed in 6 biological replicates. In total, we identified 8693 protein groups. Of these, approximately 8400 were identified per cell line, 8017 were shared between cell lines (**Figure S7A**) and a mean of approximately 7,300 protein groups was identified per run (**Figure S7B**). Principal component analysis showed that most changes happened after 24 hours of exposure to IFN-γ in all three models (**Figure S7C-E**). To check that the experiment worked as expected, we employed Single Sample Gene Set Enrichment Analysis (ssGSEA) using two IFN-γ gene signatures from the Molecular Signatures Database (MSigDB) (45). Both signatures showed a significant increase at both time points (**Figure S7F-G**). Next, we performed differential expression analysis to identify proteins significantly regulated by IFN-γ in the same direction in all models. 26 and 125 proteins were up-regulated by IFN-γ at 6 and 24 hours, respectively, while 4 and 119 proteins were downregulated at the same time points (**Figure S8A**). These proteins showed a high degree of similarity between the different patients (**Figure S8B**). We then analyzed the abundance over time of selected core IFN-γ markers (STAT1, IRF1 and CIITA) (**Figure S8C**). All proteins were detected in all models at both time points. STAT1 showed increased expression after both 2 (Log2 fold-change: 0.52; p value: 3e-16) and 6 hours of co-culture (Log2 fold-change: 2.3; p value: 4e-55). IRF1 and CIITA were consistently detected only upon IFN-γ stimulation. Since these two proteins showed a high number of missing values in the control condition, we searched for other proteins with a similar pattern, and identified 18 such proteins on the IFN-γ side and 12 on the control side (**Figure S8D**). When we compared newly synthesised proteins upon T cell attack with those affected by IFN-γ treatment (**Figure S9A**), we observed minimal overlap among down-regulated proteins, whereas up-regulated proteins were largely shared between the two conditions (**Figure 3C**). Nearly half of the proteins up-regulated by IFN-γ at 24 hours were also induced after just 6 hours of T cell attack, suggesting a synergistic response likely driven by the combined action of IFN-γ and other cytokines released during co-culture, as well as the activation of additional signaling pathways. This observation was supported by a stronger correlation between fold changes in the co-culture and those induced by IFN-γ at 24 hours (R² = 0.2; **Figure S9B**), compared to the weaker correlation observed at 6 hours (R² = 0.004; **Figure S9C**).

### Cell type-resolved analysis of protein turnover upon T cell attack

To identify proteins undergoing differential degradation upon co-culture, we excluded those regulated in the same direction in the medium-heavy channel, thereby minimizing the confounding effect of protein synthesis incorporating recycled amino acids (**Figure S10A-B**). Among the proteins differentially degraded in tumor cells (**Figure 4A**), we identified several with known roles in immune regulation. Notably, degradation of JAK2 may reflect an early immune evasion mechanism by transiently suppressing IFN-γ signaling. Consistent with this, loss-of-function mutations in JAK2 have been linked to acquired resistance to PD1 blockade immunotherapy in melanoma patients (46). Along the same line, degradation of Notch1 may help tumor cells escape T cell killing, as constitutive Notch1 signaling activity was shown to sensitize tumor cells to immunotherapy (47). Similarly, increased stability of Tyrosinase (TYR) may also contribute to early resistance to immunotherapy, as TYR has been shown to inhibit T cell anti-tumor activity in melanoma (48). Conversely, degradation of TRAF3, a negative regulator of the NFκB pathway, may support early immune activation by promoting MHC-I expression independently of PD-L1 induction (49) and by reducing TNF cytotoxicity threshold (50). On the TILs side, we also identified several immune-related proteins with altered degradation profiles (**Figure 4B**). For example, degradation of CD6 may enhance T cell activation, consistent with its known inhibitory role in immune responses (51). In contrast, degradation of the tetraspanin CD81 may dampen T cell activation and cytokine production (52).

**Figure 4.**
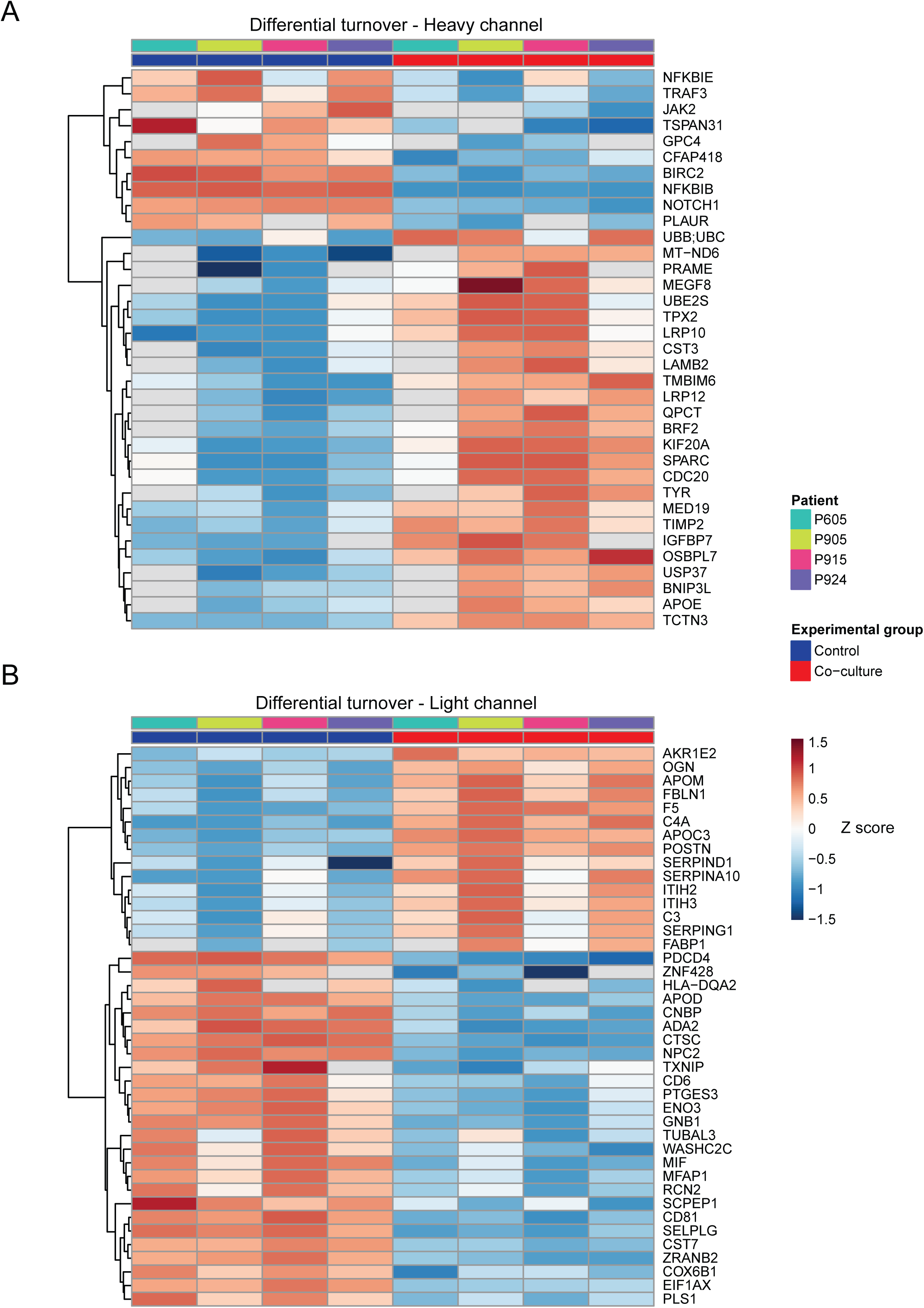
Cell type-resolved analysis of protein turnover upon T cell attack. Heatmaps of Z-scored log2 MS intensities of proteins significantly regulated in the heavy (**A**) or light (**B**) channel and not showing the same trend in the medium-heavy channel. Z-score was performed within each protein group and patient. The six replicates were averaged. Rows were clustered with euclidean distance, while the column order was fixed. The selection criteria for which proteins are included in the heatmaps are explained in Figure S10.

### Linking proteomic responses to functional outcomes of T cell attack

Whole-genome CRISPR-Cas9 screens have been widely used to dissect T cell-tumor interactions and identify genes that influence tumor cell survival during immune attack (53). In this study, we employed *in vitro* CRISPR screen data from two melanoma models: B16 cells engineered to express the model antigen ovalbumin (Ova) (39) and D10 cells that endogenously express the tumor-associated antigen MART-1 (40). In both systems, tumor cells were challenged with antigen-specific donor CD8⁺ T cells. Genes enriched at the endpoint of the screen are those whose knock-out improved cell survival under T cell pressure, while depleted genes are those whose loss reduced tumor viability. To integrate these findings with protein-level responses, we combined CRISPR screen results with our proteomic data from tumor cells exposed to T cell attack. Based on the direction of change in both datasets, we defined two groups of proteins: one where genes were either enriched and upregulated or depleted and downregulated, suggesting a potential role in promoting sensitivity to T cell killing (**Figure S11A**); and another where genes were enriched and downregulated or depleted and upregulated, indicating a possible contribution to resistance (**Figure S11B**). In the resistance group, for example, we identified BIRC2 inhibition and KDM2A, whose inhibition has been previously associated with increased sensitivity to T cell killing (47, 54). Notably, BIRC2 exhibited opposite regulation patterns between the heavy and medium-heavy channels, being downregulated (i.e., degraded) in tumor cells (**Figure 4A**), while its synthesis was increased. This may reflect a time-resolved regulatory mechanism, where an early immune response is promoted via enhanced degradation of BIRC2, followed by a delayed immune evasion strategy mediated through increased protein synthesis. We also identified several IFN-γ targets in this analysis. In the sensitivity proteins, we found TAP2, whose downregulation was previously shown to drive immune evasion and immunotherapy resistance (55). In this group, we also identified STAT1 and IRF1, two IFN-γ targets (**Figure 3C**). Notably, IFN-γ signature was previously associated with a high correlation to immune checkpoint blockade (ICB) therapy (56). However, some IFN-γ targets (MCL1, PARP12, IFI30 and OAS1) showed the opposite trend, being present in the resistance group, indicating they may contribute to the protumorigenic role of IFN-γ signaling, promoting immune evasion (57).

### CRTAM: a novel reactive cytotoxic T lymphocytes activation marker

To demonstrate that our data captures biologically meaningful changes, we functionally validated the role of the transmembrane protein Cytotoxic And Regulatory T Cell Molecule (CRTAM) during T cell attack of melanoma cells. CRTAM was uniquely detected in the medium-heavy channel of the proteomics data after 6 hours of co-culture in three out of four patients (**Figure 3B** and **S12A**). Flow cytometry analysis across all four patients confirmed CRTAM upregulation almost exclusively on CD8^+^ cytotoxic T lymphocytes (CTLs) following tumor engagement (**Figure 5A** and **Figure S12B-C**), with P905 and P915 showing more than 15% CRTAM⁺ CD8⁺ TILs (**Figure S12B**). This upregulation required CTL activation through the TCR/CD3 complex, was further enhanced by CD28 co-stimulation, and was independent of IFN-γ (**Figure 5A-B**). CRTAM expression was rapidly induced upon co-culture, reaching a maximum level between 7 and 24 hours before returning to baseline by 48 hours (Figure **S12D**), making CRTAM an early CTL activation marker. CRTAM-positive CTLs co-expressed the reactivity marker TNFRSF9/CD137 and the survival marker IL2RA/CD25 (**Figure 5C** and **S13A**), and showed increased secretion of IFN-γ compared to CRTAM-negative CTLs (**Figure 5D** and **Figure S13B**). This suggests that CRTAM identifies a subset of tumor-infiltrating CTLs with enhanced anti-tumor activity. In a single-cell RNA sequencing (scRNA-seq) dataset from 33 melanoma tumors (41), CRTAM expression was confirmed predominantly on CTLs (**Figure S14A**), both CD137 and IFN-γ transcripts were significantly enriched in CRTAM-high CTLs (**Figure S14B-E**), while IL2RG/CD132 was enriched in CRTAM-positive cells (**Figure S14F**).

**Figure 5.**
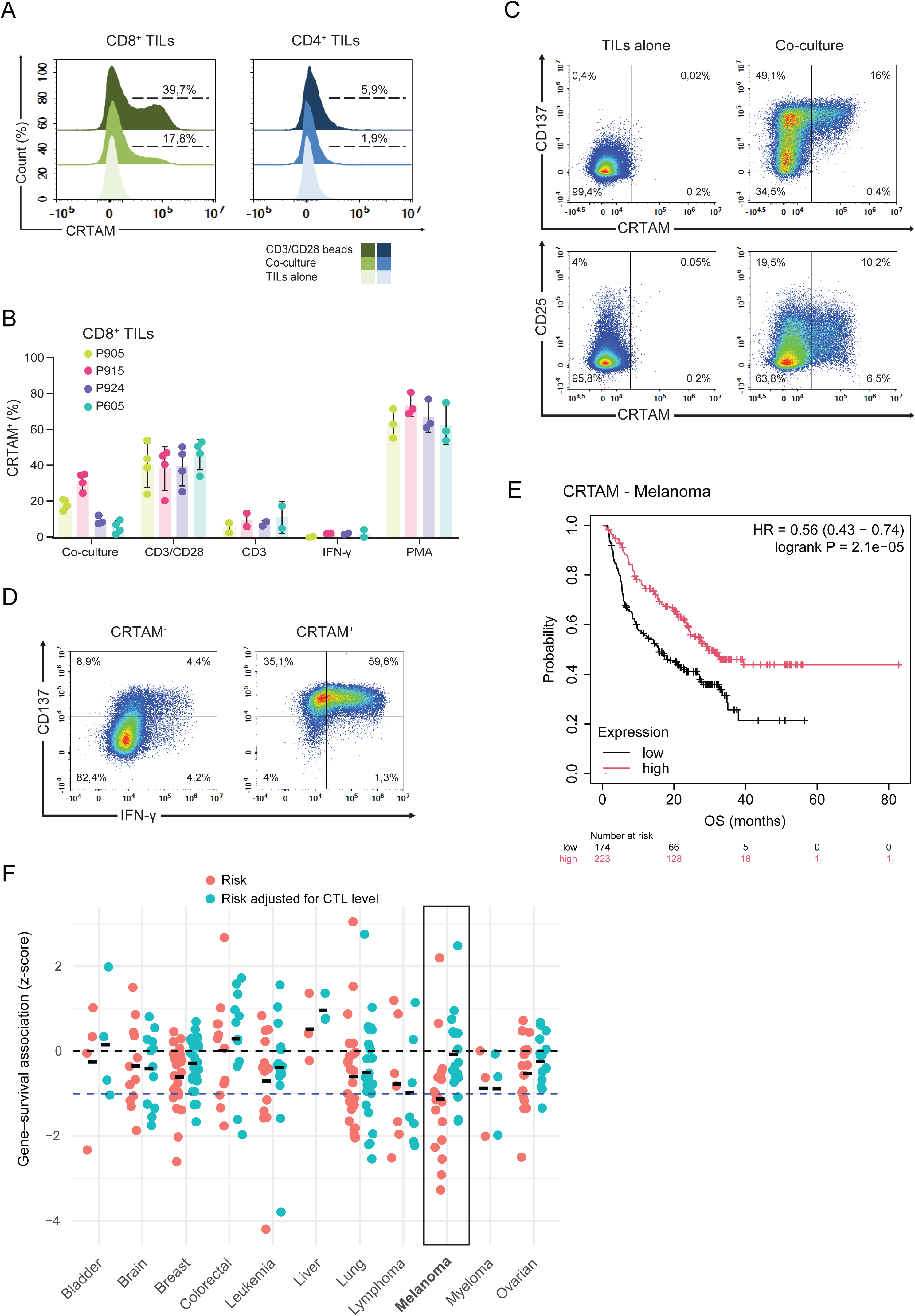
Functional role of CRTAM in melanoma immune recognition. **A**. CRTAM surface expression on CD4⁺ and CD8⁺ TILs from patient 905, assessed by flow cytometry after 6 hours of stimulation with CD3/CD28 beads or co-culture with autologous melanoma cells at an E:T ratio of 1:1. **B**. Percentage of CRTAM in CD8⁺ TILs from all four patients after 6 hours of co-culture with autologous melanoma cells at an E:T ratio of 1:1, or stimulation with anti-CD3 antibody, CD3/CD28 beads, IFN-γ, and PMA. Data are presented as mean ± SD (n = 2-4 biological replicates, each with three technical replicates). **C**. Representative density plots of flow cytometric analysis showing surface co-expression of CRTAM with CD137 or CD25 on CD8⁺ TILs from patient 905 after 6 hours of co-culture with autologous melanoma cells at an E:T ratio of 1:1. **D**. Intracellular expression by flow cytometry of CD137 and IFN-γ in CRTAM⁺ and CRTAM⁻ CD8⁺ TILs from patient 915 after 8 hours of co-culture with autologous melanoma cells at an E:T ratio of 1:1. **E**. KM curve showing the association between CRTAM gene expression and survival in a cohort of 423 melanoma patients undergoing treatment with immune checkpoint inhibitors. **F**. Z-score of the effect of CRTAM expression on overall survival across multiple cancer types in a CoxPH model, before and after adjusting for CTL cell infiltration. A negative z-score means that higher CRTAM expression is associated with lower death risk. Melanoma cohorts are circled in blue.

To assess the clinical relevance of CRTAM in CTL-mediated melanoma killing, we analyzed the correlation between CRTAM gene expression and overall survival (OS) in a clinical cohort of 423 patients with melanoma undergoing treatment with immune checkpoints inhibitors (58), and observed that high CRTAM expression predicted improved OS (**Figure 5E**), suggesting that CRTAM expression may serve as a surrogate marker of anti-tumor immune activity and clinical benefit. To assess whether the association between CRTAM and improved OS extended beyond this cohort and melanoma, we evaluated the prognostic effect of CRTAM expression across multiple melanoma cohorts and cancer types using a Cox proportional hazards (CoxPH) regression model implemented in the TIDE (Tumor Immune Dysfunction and Exclusion) algorithm (59, 60). High CRTAM expression was associated with reduced risk of death in most cancer types (all except bladder, colorectal, and liver cancer), with melanoma showing the strongest and most consistent protective effect (**Figure 5F**). Among 19 melanoma cohorts, only two did not show a survival benefit (**Figure S15A**). Adjusting for CTL infiltration markedly reduced the predictive value of CRTAM in melanoma, indicating that its association with survival was largely mediated by CTL abundance (**Figure 5F** and **S15A**). In contrast, in other cancers, such as lymphoma and myeloma, CRTAM retained prognostic power after adjusting for CTL infiltration (**Figure 5F**). Interestingly, in these cancer types, CD8A, the canonical marker of CTL infiltration (61), showed no prognostic power (**Figure S15B**). Consistent with this, CRTAM expression positively correlated with CTL levels across cohorts, most strongly in melanoma, but consistently less robustly than CD8A (**Figure S15C-D**). Together, these findings suggest that CRTAM is not a general proxy for CTL infiltration, but rather reflects a subset of CTLs whose presence is associated with favorable clinical outcome, consistent with a role in effective anti-tumor immunity.

### Cell type-resolved analysis of protein phosphorylation upon T cell attack

On the phosphoproteome data, we performed differential expression analysis separately for each SILAC channel to identify phosphorylation sites consistently regulated upon T cell attack across all models. Since peptides containing medium-heavy-labeled amino acids could originate from either tumor cells or TILs, we focused our analysis on the heavy (tumor-derived) and light (TIL-derived) channels. We identified 1145 up-regulated by T cell attack and 153 down-regulated phosphosites in the heavy channel, and 743 up-regulated and 128 down-regulated phosphosites in the light channel (**Figure S16A**). Phosphorylation patterns were profoundly different between melanoma cells and TILs, (**Figure 6A**). Among phosphosites with an associated function in the PhosphoSitePlus database (62) and identified in both cell types, the site most upregulated preferentially in TILs was serine 39 on Vimentin (**Figure 6B**), an AKT1 target known to affect cell motility and cytoskeleton reorganization . Among the most regulated functional phosphosites in TILs but not identified in melanoma cells, we found several T cell activation markers, for example tyrosine 142 on CD3ζ (**Figure 6C**), a crucial component of the T cell receptor (TCR) signaling complex. The most regulated functional phosphosite in melanoma cells, also identified in TILs, was serine 2612 on DNA-PK (**Figure 6D**), known to enhance the process of DNA repair. Among the sites commonly regulated in both cell types, we identified serine 320 on RIPK1 (**Figure 6E**), a target of MAPKAPK2 (MK2) and TAK1 kinases in response to inflammatory stimuli such as TNF-α. Phosphorylation at this site inhibits RIPK1-mediated apoptosis by preventing its interaction with FADD and caspase-8 (63–66).

**Figure 6.**
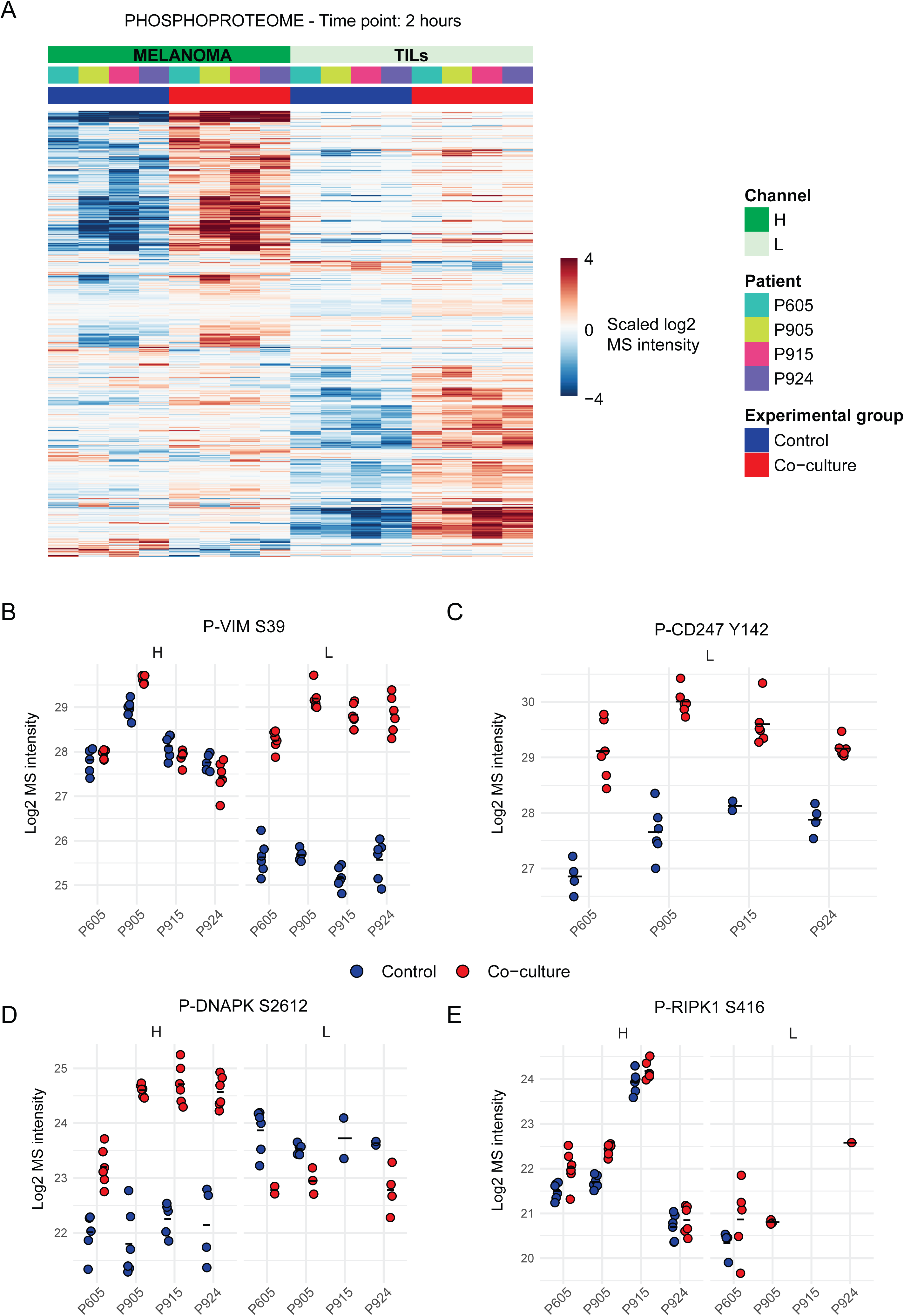
Cell type-resolved analysis of protein phosphorylation upon T cell attack. **A**. Heatmap of scaled log2 MS intensities of the 2169 significantly regulated phosphosites in the heavy and light channels. Missing values were imputed with the minimum value of the channel-specific filtered matrix before scaling. Scaling was performed within each protein group, patient and channel. The six replicates were averaged after scaling. Phosphosites uniquely present in one channel were imputed with 0s after scaling. Rows were clustered with maximum metric distance, while the column order was fixed. **B-E**. Log2 MS intensities of selected phosphosites. The black dash represents the mean.

Using pathway enrichment analysis, in melanoma cells, we observed enrichment of DNA damage response (DDR) pathways under T-cell attack, including those mediated by ATM and ATR signaling (**Figure 7A**). These findings corroborated our previous analyses. Interestingly, melanoma cells also exhibited enrichment in innate immune signaling pathways typically associated with antiviral and antibacterial responses (RIG-I-like receptor signaling pathway), suggesting that the intracellular signaling landscape of tumor cells under immune attack mirrors the response of immune cells to pathogenic insults. In contrast, several pathways involved in cell motility were significantly downregulated upon attack, potentially reflecting a shift from a migratory to a defensive state. Conversely, in TILs, we observed significant enrichment of pathways related to cell motility and cytoskeletal remodeling upon engagement with melanoma cells, consistent with active migration and immune synapse formation. Additionally, pathways associated with T-cell receptor (TCR) activation were strongly enriched, confirming that TILs were functionally responding to tumor antigens during the interaction. Very few pathways showed a shared regulation between the two cell types. This was confirmed by a statistically significant negative linear association between the two cell types at pathway level (intercept = -0.16, p = 0.007), with the model explaining only a small fraction of the variance (R² = 0.03), indicating that the overall pathway patterns are mostly unrelated in the two cell types (**Figure S16B**).

**Figure 7.**
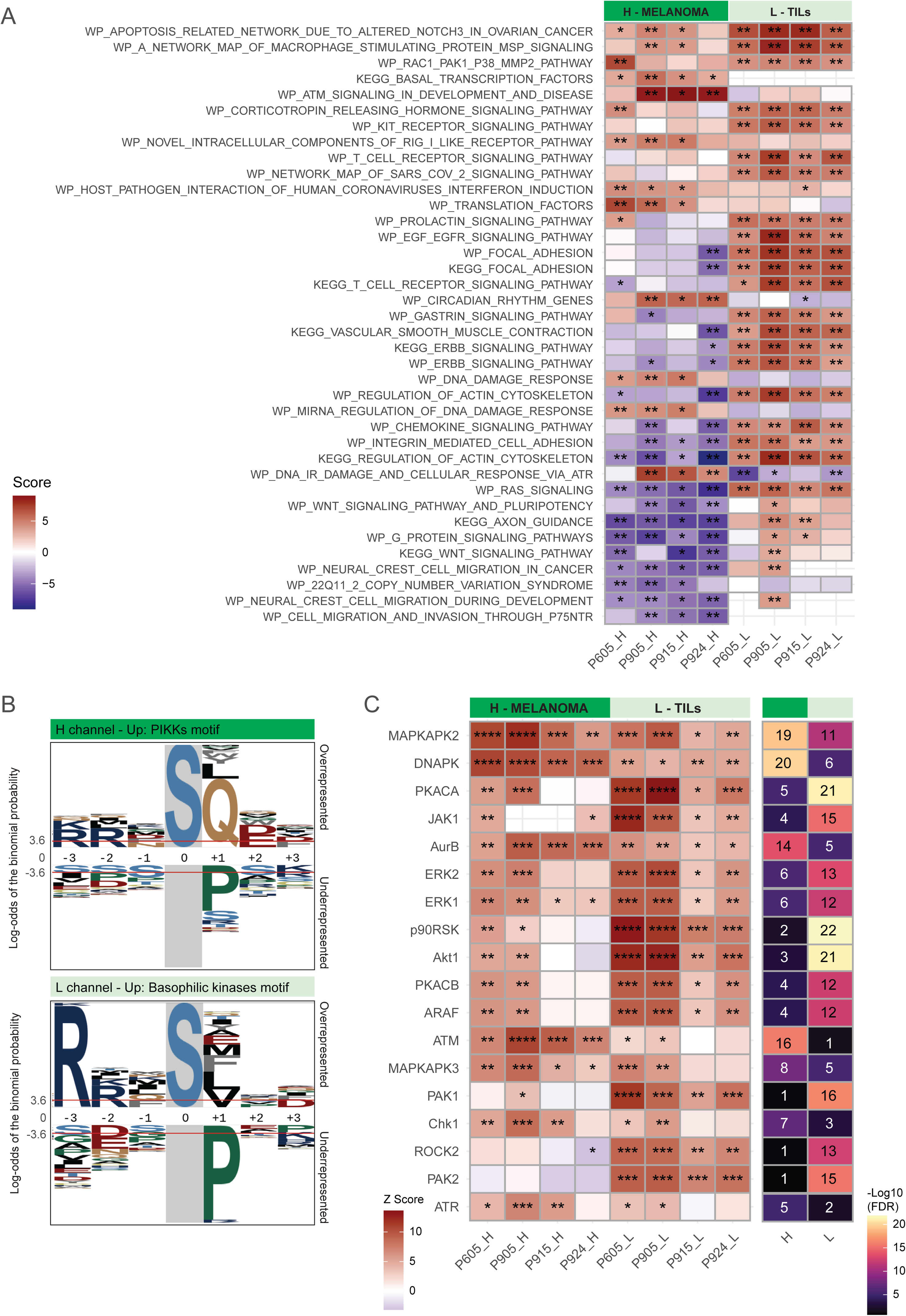
Cell type-resolved analysis of phosphorylation-and kinase-driven signaling upon T cell attack. **A**. Heatmap of gene-centric-redundant ssGSEA scores. The 20 pathways with the highest sum of -log10(adjusted p) across the four patients per channel are shown, for a total of 38 unique pathways. Significance is represented by asterisks: * = adjusted p <= 0.05; ** = adjusted p <= 0.01; *** = adjusted p <= 0.001. **B**. Amino acid motifs over-represented among up-regulated phosphosites in each channel. **C**. Heatmaps of RoKAI Z-scores (left) and -log10(FDR) (right). Only kinases expressed on the proteome level are shown. The 10 kinases with the highest sum of -log10(FDR) across the four patients per channel were selected for this plot, for a total of 18 unique kinases. Significance is represented by asterisks: * = FDR <= 0.05; ** = FDR <= 1e-5; *** = FDR <= 1e-10; **** = FDR <= 1e-20.

In melanoma cells, motif enrichment analysis (32) highlighted the over-representation of a glutamine residue at the +1 position upon attack, which represents the sequence motif for the phosphatidylinositol 3-kinase-related kinases (PIKKs): ATM, ATR and DNA-PK (67, 68) (**Figure 7B**). Conversely In TILs, the basophilic kinase motif R/K-R/K-x-pS/pT was over-represented upon interaction with melanoma cells, highlighting major differences in kinase activation between the two cell types.

Next, we employed the RoKAI algorithm to assess kinase activity upon T cell attack, by analyzing alterations in the phosphorylation of known kinase substrates and their functional network (33). Among the kinases preferentially enriched in melanoma cells upon attack we identified DNA-PK, Aurora Kinase B, ATM, Chk1 and ATR (**Figure 7C**), confirming the former analyses. Most of these kinases are involved in DDR: DNA-PK and ATM are involved in DNA repair following double strand breaks (DSBs) through non-homologous end joining (NHEJ) and Homologous Recombination (HR), respectively (69); ATR is the master regulator of the response to replication stress and single-stranded DNA (69), while Chk1 acts downstream ATR (70). MAPKAPK2 and MAPKAPK3 were also preferentially enriched in tumor cells. These are stress-responsive protein kinases regulated through direct phosphorylation by p38 MAPK, involved in several cellular processes including stress and inflammatory responses, cytokine production and DNA damage response (71).

Multiple kinases preferentially enriched in TILs were basophilic kinases (PKACA, p90RSK, Akt1, PKACB, PAK1, ROCK2, PAK2), confirming the motif analysis. Moreover, we identified several members of the canonical MAPK signaling pathway, including ERK1 and ERK2, and their downstream target p90RSK. Additionally, both PAK1 and PAK2 showed increased activity, confirming previous findings by the Stecker’s lab (10). Akt1 was also preferentially activated in TILs, as well as the protein Kinase cAMP-activated (PRKACA and PRKACB). ROCK2 was also markedly enriched in TILS, and it is known to control cellular processes related to the cytoskeleton and cell motility, explaining the enrichment of cytoskeleton-related pathways in TILs (**Figure 7A**). To confirm the observed kinase activation pattern, we analyzed kinase activity using PTM-Signature Enrichment Analysis (PTM-SEA). This analysis confirmed the preferential enrichment of MAPKAPK2, AURKB, MAPKAPK3, Chk1 and ATM in melanoma cells (**Figure S16C**). Additionally, it showed preferential activation of all four p38 MAP kinases (MAPK11, 12, 13 and 14), which are upstream of MAPKAPK2 and MAPKAPK3. This analysis also confirmed preferential enrichment in TILs of PRKACA, Akt and p90RSK.

All in all, these analyses confirmed the ability of our approach to resolve phosphorylation-specific signaling events with cell-type resolution.

### DNA-PK: a potential immune resistance kinase in melanoma

Activation of DDR kinases was prominent in our data. Several phosphorylation sites on the DNA-PK kinase were up-regulated on melanoma cells upon interaction with TILs, many of which are known to promote DNA repair (**Figure S17A-B**).

To investigate the functional relevance of DNA-PK during T cell attack, we first analyzed the association between DNA-PK gene expression and OS in a clinical cohort of patients treated with immune checkpoints inhibitors (48), and found that high DNA-PK expression predicted worse OS (**Figure S17C**). We confirmed this negative prognostic association across multiple cancer types (**Figure S17D**). Importantly, the predictive value of DNA-PK remained after adjusting for the level of T-cell infiltration. In line with this, DNA-PK gene expression negatively correlated only weakly with CTL infiltration (**Figure S17E**), suggesting that DNA-PK expression influences survival independently of T-cell abundance in the TME.

To further dissect the role of DNA-PK in tumor immune evasion, we examined data from the negative CRISPR screen by Zhang et al. (40) for DNA-PK, ATM and key immune checkpoints (PD-L1, PD-L2, TIM-3 and IDO1). None of these genes showed strong negative selection, meaning that their knockout did not significantly impair melanoma cell survival (**Figure S17F**). Consistently, treatment with DNA-PK inhibitors, alone or in combination with ATM inhibitors, did not alter tumor cell killing nor did it affect IFNγ-mediated cytotoxicity in our system (data not shown). However, while DNA-PK knockout did not reach high statistical significance, it showed a negative rank close to the best-performing immune checkpoints, highlighting its potential role as an immune resistance factor *in vivo*, suggesting the role of this kinase in melanoma immune escape.

## Discussion

In this study, we established a SILAC-DIA phosphoproteomics workflow to resolve early, cell type-specific signaling events during interactions between patient-derived melanoma cells and matched autologous TILs. By combining triple SILAC labeling with Orbitrap Astral-based DIA, we overcame a major barrier in proteomic studies of co-cultures, that is the loss of cell identity upon lysis, while avoiding the need for physical separation of cell types by FACS. This strategy enabled a deep and quantitative analysis of phosphorylation site dynamics, protein turnover, and the newly synthesized proteome from both tumor and immune compartments within the same experiment. Our approach revealed coordinated yet distinct responses in melanoma cells and TILs during the early stages of T cell-mediated attack.

In the immune compartment, we identified the cell surface receptor CRTAM (Class I-restricted T cell-associated molecule, also known as CD355) as a marker of a tumor-reactive CTL subpopulation. Consistent with previous studies, CRTAM expression was rapidly induced upon CTL stimulation and promoted secretion of key cytokines such as IFN-γ (72, 73). Clinically, CRTAM could be explored as a biomarker to stratify melanoma patients by the presence and activity of tumor-reactive T cells, aiding prediction of immune checkpoint blockade responses or monitoring efficacy of adoptive T cell therapies, including TIL-based approaches. Moreover, CRTAM itself, or pathways regulating its expression, may represent actionable targets to enhance CTL cytotoxicity in the tumor microenvironment.

In the tumor compartment, we observed rapid activation of kinases linked to the DNA damage response, including DNA-dependent protein kinase (DNA-PK/PRKDC). While DNA-PK is best known for its role in non-homologous end joining during DNA repair (74), recent studies show it also modulates tumor-immune interactions. Inhibiting DNA-PK with NU7441 increased MHC-I expression, reduced immunosuppressive proteins, expanded the tumor neoantigen landscape, and promoted antigen presentation in melanoma models (75, 76). Our detection of DNA-PK activation within minutes of T cell engagement suggests it is part of an immediate adaptive response to immune attack. Together with published data, these results support targeting DNA-PK as a strategy to impair tumor survival pathways and enhance tumor immunogenicity, potentially creating a therapeutic window for combination with immunotherapies.

Our model system, a 2D co-culture of patient-derived melanoma cells and autologous TILs, provides a simple and controllable platform to dissect direct tumor-T cell interactions, but it has several limitations. First, it lacks other immune cell populations, three-dimensional architecture, extracellular matrix, and stromal components such as fibroblasts, all of which can shape immune responses. In line with this, several clinically relevant immune checkpoint pathways showed little or no functional effect in this system (**Figure S17E**), highlighting the need to validate candidate biomarkers and targets, such as DNA-PK, in more physiologically relevant models, including organoid-based co-cultures or humanized mouse systems. Second, newly synthesized proteins during co-culture are incorporated into the medium-heavy SILAC channel without retaining information about the cell type of origin, meaning cell-type specificity is preserved only for pre-existing proteins. This introduces a time-dependent constraint: as co-culture progresses and more proteins are synthesized, a larger fraction of the proteome shifts into the medium-heavy channel, progressively reducing the ability to assign them to a specific cell type. Third, differences in cell size and protein content result in lower proteomic depth for smaller immune cells compared to melanoma cells. Fourth, triple SILAC labeling limits the approach to studying two cell types, as all three isotopic states are required to distinguish compartments and track new protein synthesis. Despite these limitations, our work establishes a broadly applicable experimental and analytical framework for dissecting cell type-specific signaling in intact mixed-cell systems without physical separation. This approach is particularly suited to studying dynamic interactions in heterogeneous microenvironments where conventional isolation strategies may disrupt labile PTMs or introduce technical bias. By enabling high-resolution mapping of early signaling events in both tumor and immune compartments, our workflow opens new opportunities for identifying therapeutic targets and biomarkers in cancer immunology. With further refinements, including integration of emerging multiplexing strategies, this methodology could be extended to more complex and clinically relevant models, ultimately bridging the gap between mechanistic discovery and translational application.

## Data Availability

The datasets generated and analyzed during the current study are available from the corresponding author upon request, subject to a collaboration agreement.

## Acknowledgments

We are grateful to all patients who donated their samples for this work. We also thank Aishwarya Gokuldass for her participation in the early stages of this project, Anne-Christine Kiel Rasmussen for performing the melanoma flow cytometry panel experiments, and Ulises Hernández Guzmán for helping with MS analysis of SILAC-labeled samples.

## Authors’ contribution

GF and JVO designed the study. GF and AWPJ optimized the SILAC DIA methodology, wrote the original draft of the manuscript and generated the figures. GF, AWPJ and IP performed experiments. GF, AWPJ and AMV analyzed data. GF generated the R code used to analyze the data. JVO and MD provided resources and coordinated the project. GF, MD and JVO supervised the project and acquired funds. All co-authors read and edited the manuscript.

---

**Figure S1.**
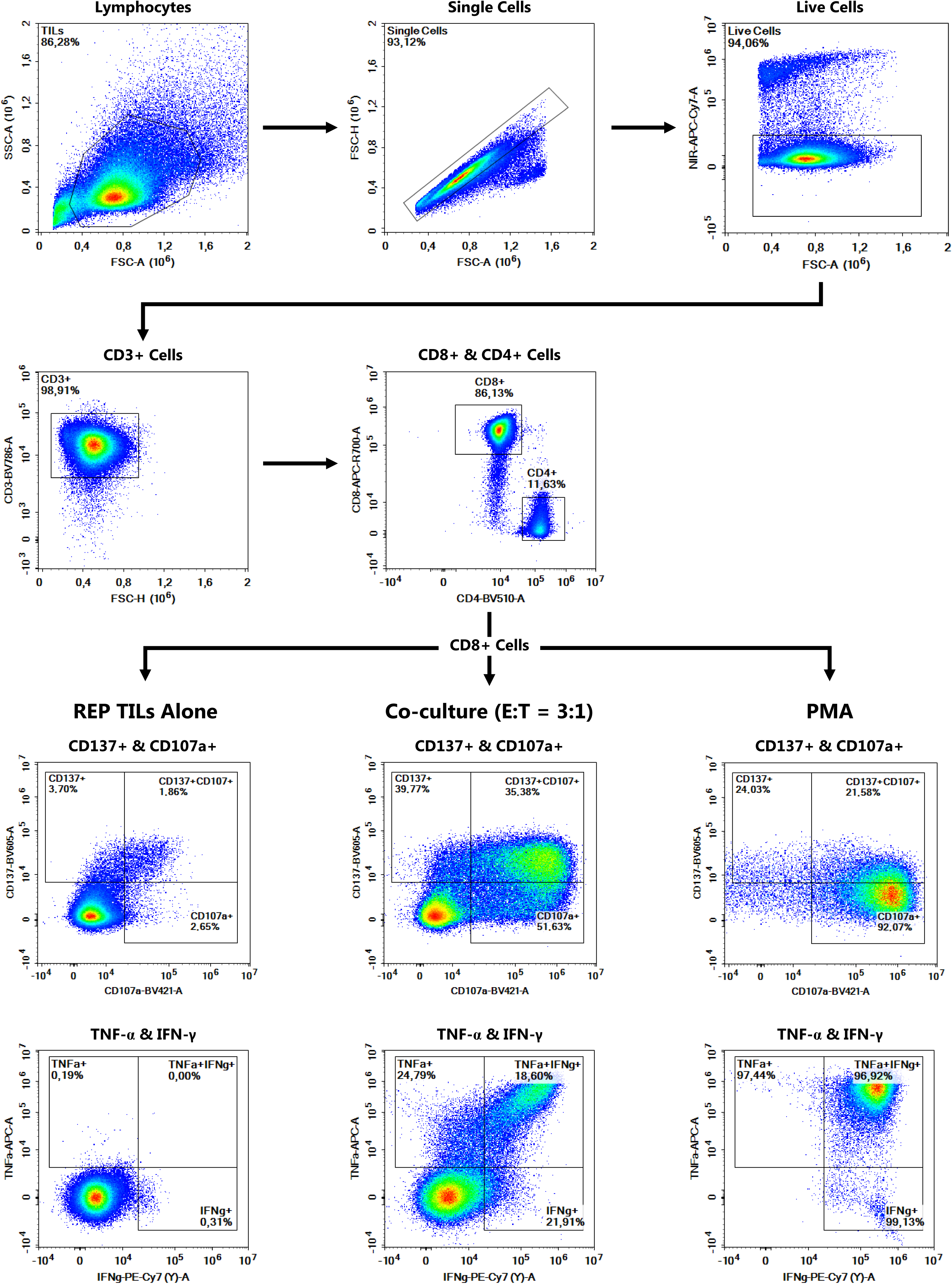
Flow cytometry gating strategy for the experiment displayed in Figure 1A. The same strategy has been used for all flow cytometry experiments included in this study.

**Figure S2.**
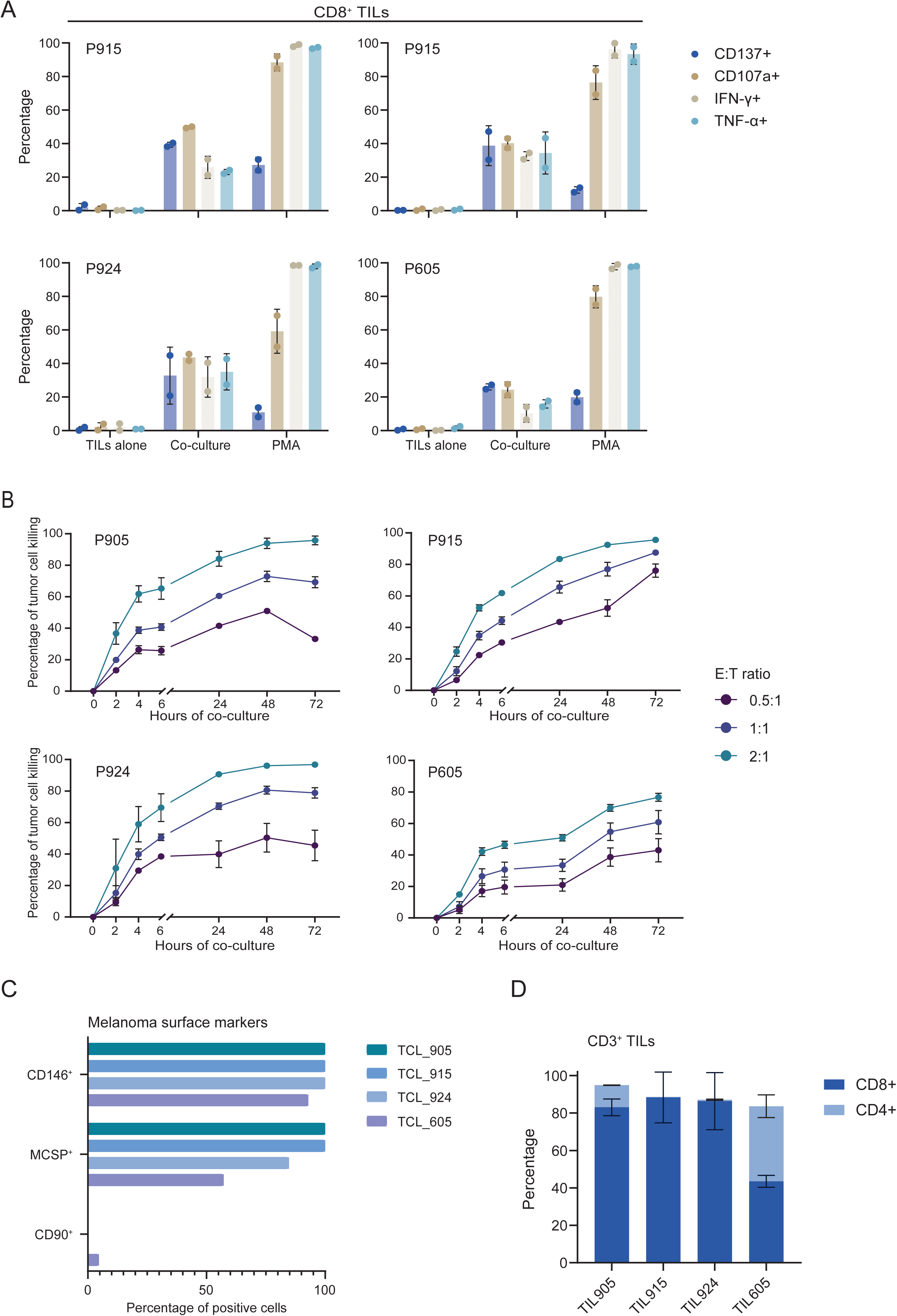
Functional characterization of tumor-infiltrating lymphocytes (TILs) and melanoma patient-derived tumor cell lines (TCLs). Related to Figure 1A-B. **A**. Flow cytometric analysis of tumor-specific CD8⁺ TIL reactivity based on upregulation of activation markers (CD137, CD107a) and intracellular cytokines (IFN-γ, TNF-α) after 8 hours of co-culture with autologous melanoma cells at an E:T ratio of 3:1. TILs alone and PMA/ionomycin stimulation served as negative and positive controls, respectively. Data are presented as mean ± SD (n=2 biological replicates). **B**. Autologous TIL-mediated killing of melanoma cells from four patients was assessed using xCELLigence real-time cell analysis at three different E:T ratios. The percentage of tumor cell killing was calculated as the Normalized Cell Index (NCI) of the tumor in co-culture divided by the NCI of tumor alone x 100. Data are shown as mean ± SD (n=3-4 technical replicates). **C**. Surface expression of two melanoma markers (CD146 and MCSP) and one fibroblast marker (CD90) in four different melanoma patient-derived TCLs. **D**. Surface expression of CD4 and CD8 by flow cytometry in CD3^+^ TILs. Data are presented as mean ± SD (n=2 biological replicates).

**Figure S3.**
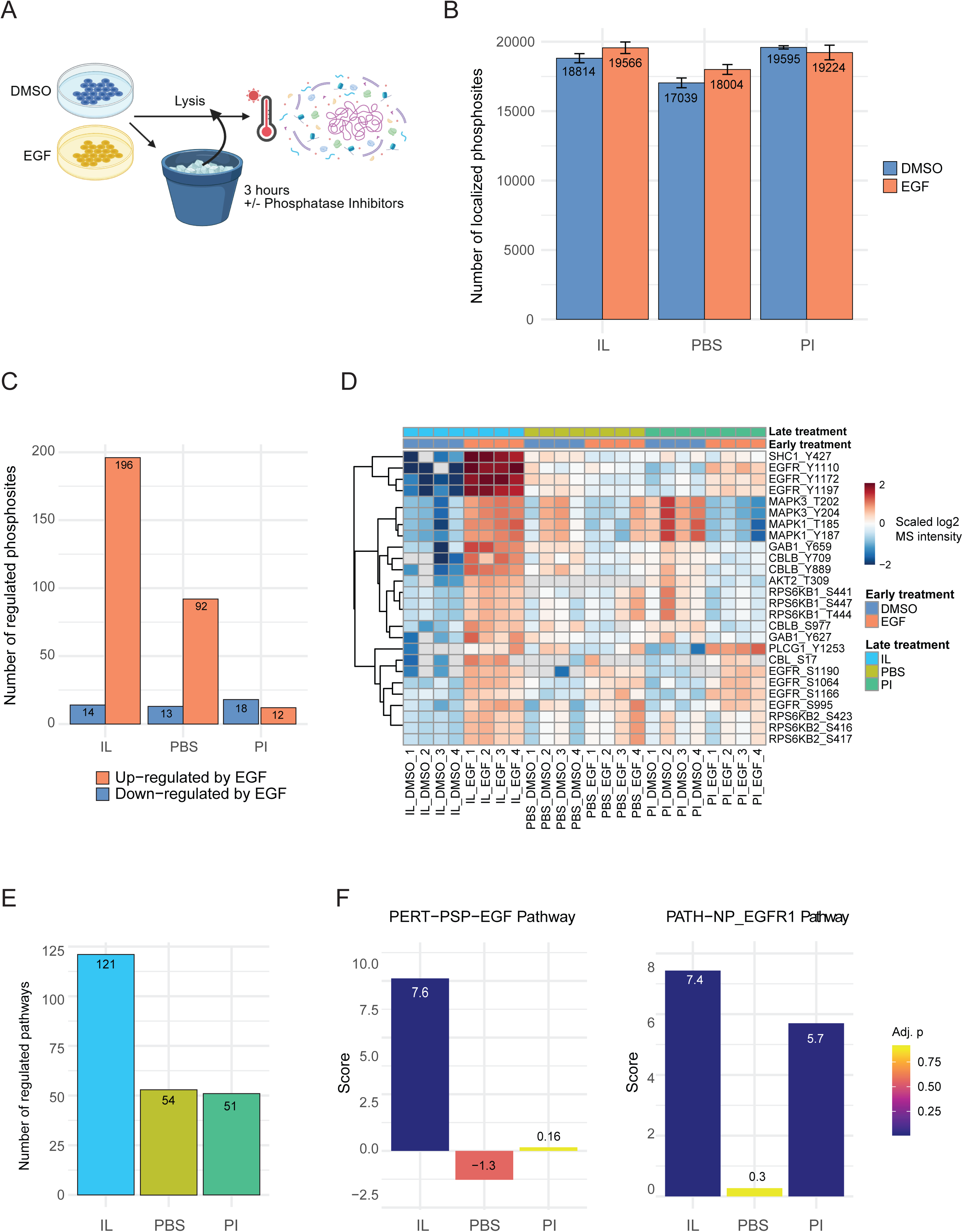
Phosphoproteomic profiling of SCC-25 cells stimulated with EGF and lysed immediately or after incubation on ice. **A**. Schematic drawing of the phosphoproteomics experiment performed in SCC-25 squamous cell carcinoma cells. Following stimulation with EGF for 8 minutes, SCC-25 were either lysed immediately using a hot lysis buffer (IL), or resuspended in PBS and further incubated on ice for three hours prior to lysis with (PI) or without (PBS) the addition of phosphatase inhibitors. Created with Biorender. **B**. Localized phosphosite identifications (localization probability >= 0.75) per condition. Data are presented as mean ± SD (n=4 biological replicates). Samples were analyzed on an 80 sample-per-day gradient. **C**. Number of up- and down-regulated phosphosites per contrast. **D**. Heatmap of scaled log2 MS intensities of the significantly regulated phosphosites in at least one contrast belonging to the ErbB KEGG signaling pathway (n=33 phosphosites). Rows were clustered with euclidean distance, while the column order was fixed. **E**. Number of significantly regulated (adjusted p <= 0.05) pathways by PTM-SEA in at least one contrast **F**. PTM-SEA score representing phosphorylation alterations in response to EGF.

**Figure S4.**
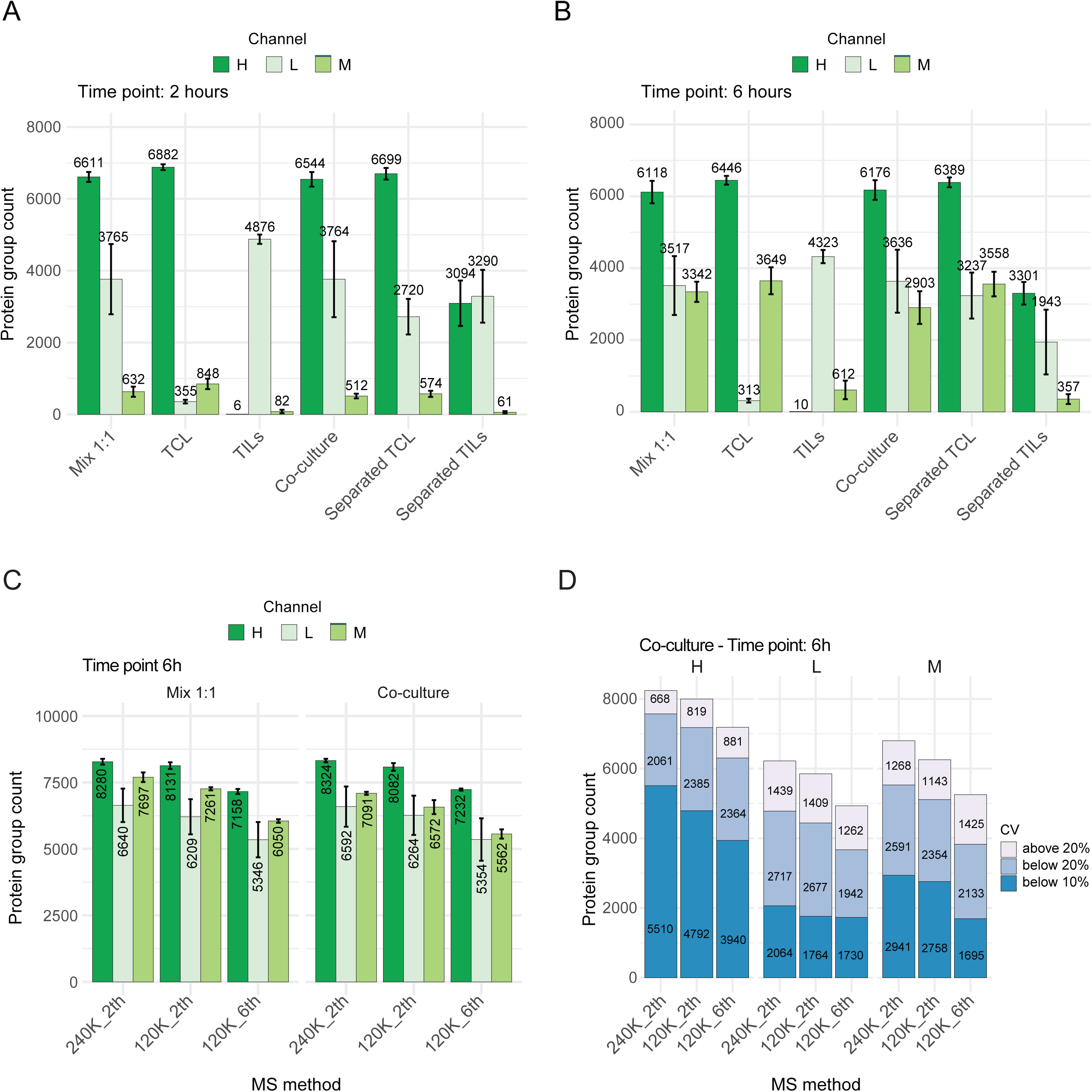
SILAC-DIA method optimization. Related to Figure 1C-D. **A-B**. Protein group identifications per channel and condition at time point 2-hour (A) and 6-hour (B) of co-culture. Data are presented as mean ± SD (n=4 biological replicates). Samples were analyzed on an 180 sample-per-day gradient. **C**. Protein group identifications per channel and MS method at time point 6-hour of co-culture. Data are presented as mean ± SD (n=4 biological replicates). Samples were analyzed on a 36 sample-per-day gradient. **D**. Protein group coefficient of variation (CV) calculated for proteins with at least 3 out of 4 valid values in the co-culture condition at time point 6-hour. Samples were analyzed on a 36 sample-per-day gradient.

**Figure S5.**
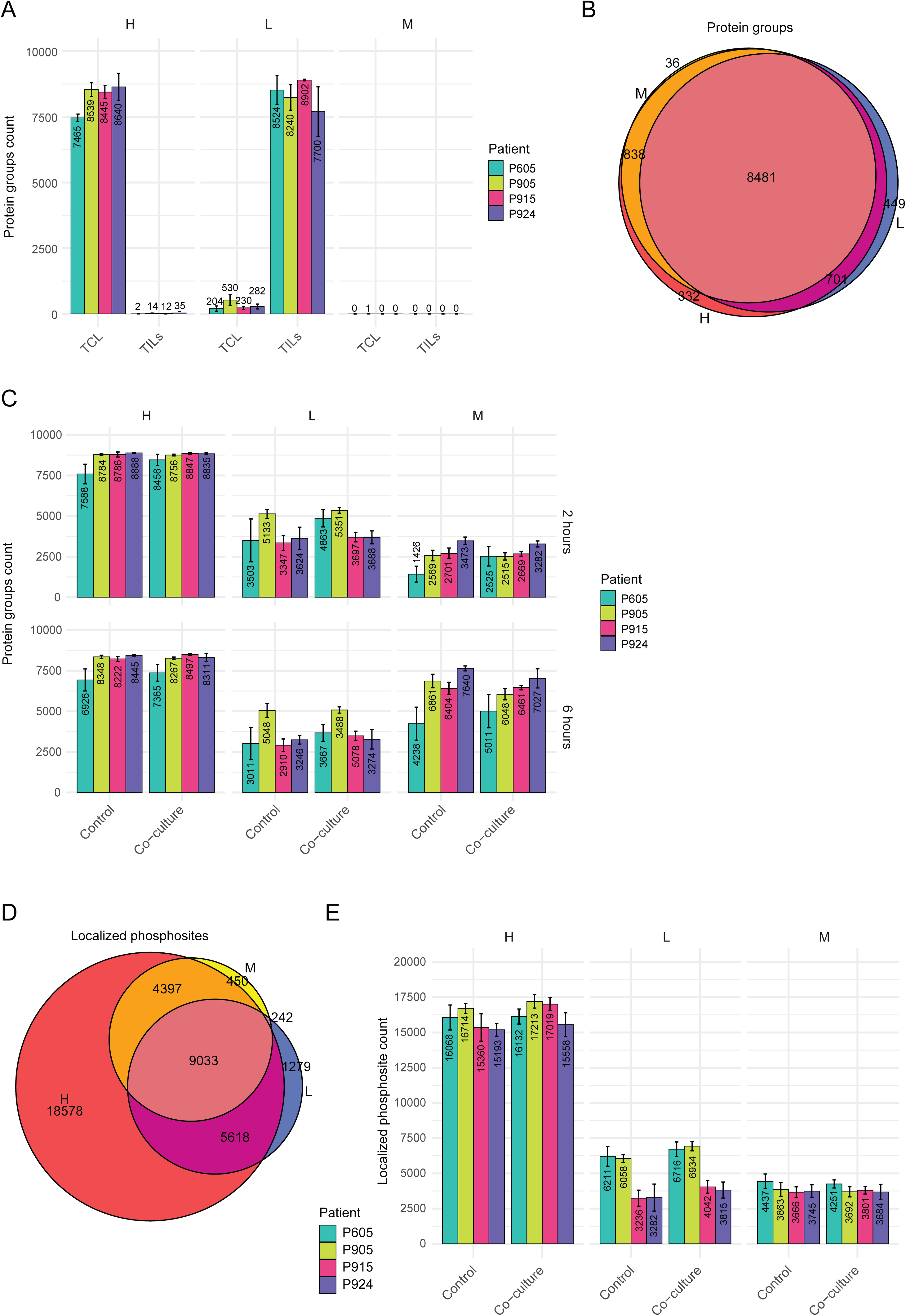
Protein and phosphosite coverage achieved in SILAC-DIA co-cultures. Related to Figure 2. **A**. Protein group identifications per channel, cell type and patient at time point 0-hours (before co-culture). Data are presented as mean ± SD (n=6 biological replicates). Samples were analyzed on a 30 sample-per-day gradient. **B**. Euler diagram showing the overlap between protein group identifications in the three different channels. **C**. Protein group identifications per channel, cell type and patient at time point 2- and 6-hours of co-culture. Data are presented as mean ± SD (n=6 biological replicates). **D**. Euler diagram showing the overlap between phosphosite identifications in the three different channels. **E**. Phosphosite identifications per channel, cell type and patient at time point 2- and 6-hours of co-culture. Data are presented as mean ± SD (n=6 biological replicates).

**Figure S6.**
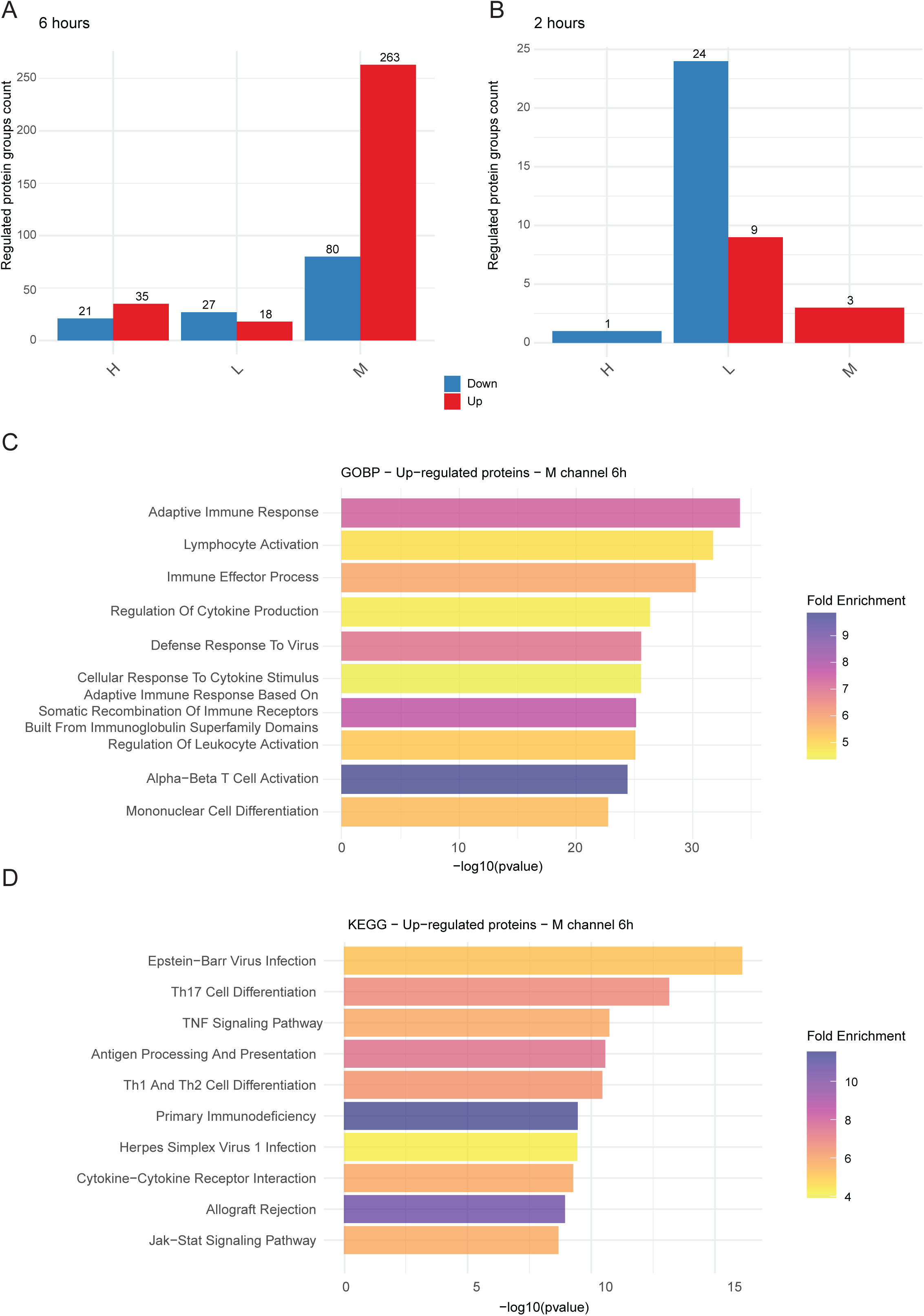
Differential protein regulation and pathway enrichment upon co-culture. Related to Figure 3A-B. **A-B**. Number of up- and down-regulated protein groups per contrast and channel at time point 6-hour (A) and 2-hour of co-culture. **C-D**. Gene ontology biological process (GOBP; C) and KEGG (D) over-representation analysis for proteins significantly regulated in the medium-heavy channel upon co-culture. These include both significantly regulated proteins and those exclusively present in one condition, for a total of 299 entries for GOBP and 301 for KEGG. The top most significant (lowest adjusted p value) terms are shown. For GOBP, redundant terms were manually removed. Fold enrichment is calculated by dividing the proportion of the proteins in a pathway by the proportion of all proteins in the background within that same pathway.

**Figure S7.**
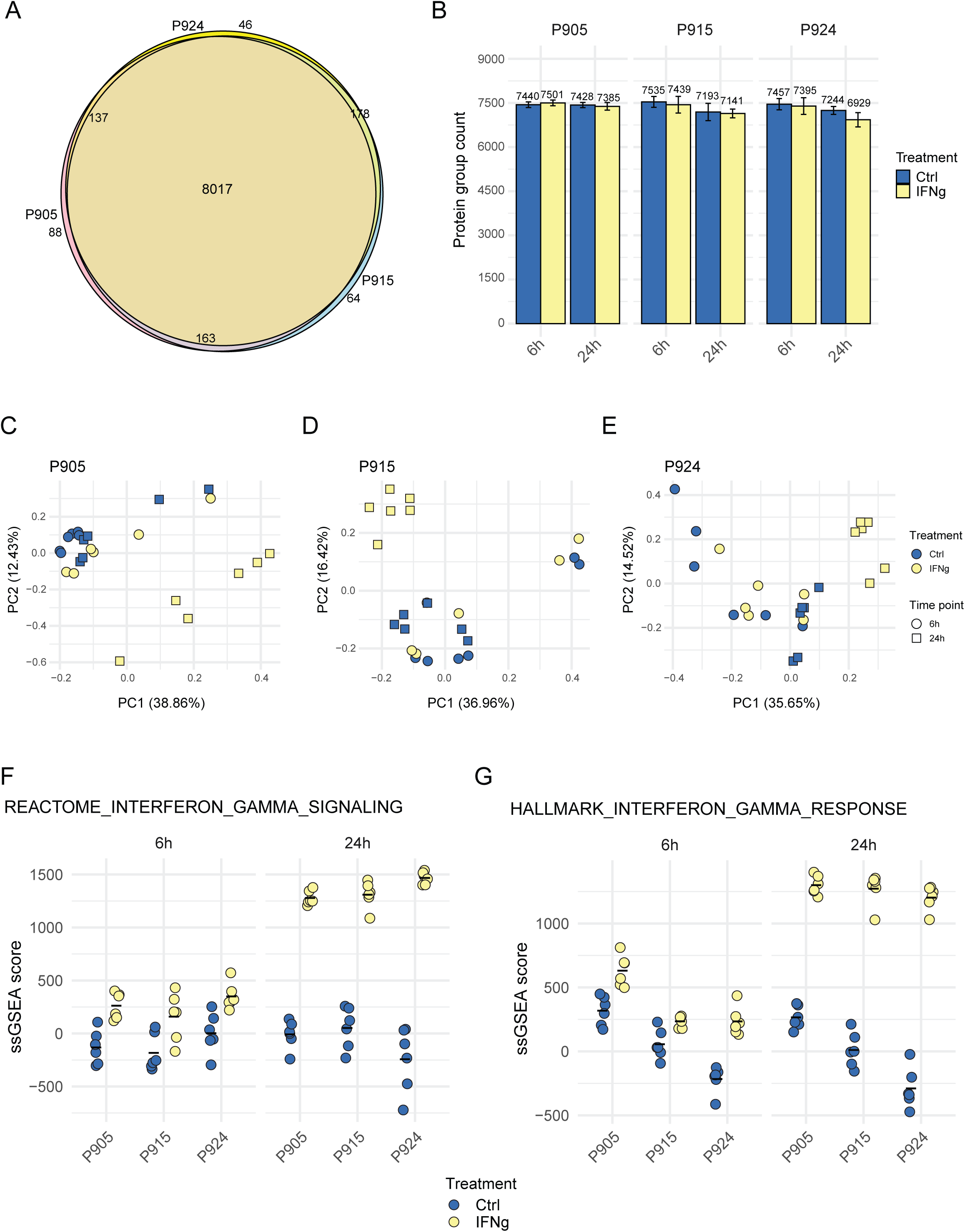
Proteome changes induced by IFN-γ in three patient-derived melanoma cell lines. **A**. Euler diagram showing the overlap between protein group identifications in the three different patients. **B**. Protein group identifications per patient and time point. Data are presented as mean ± SD (n=6 biological replicates). Samples were analyzed on an 180 sample-per-day gradient. **C-E**. Principal component analysis by patient. F-G. Single-sample gene set enrichment analysis (ssGSEA) using the REACTOME and KEGG IFN-γ signaling pathways.

**Figure S8.**
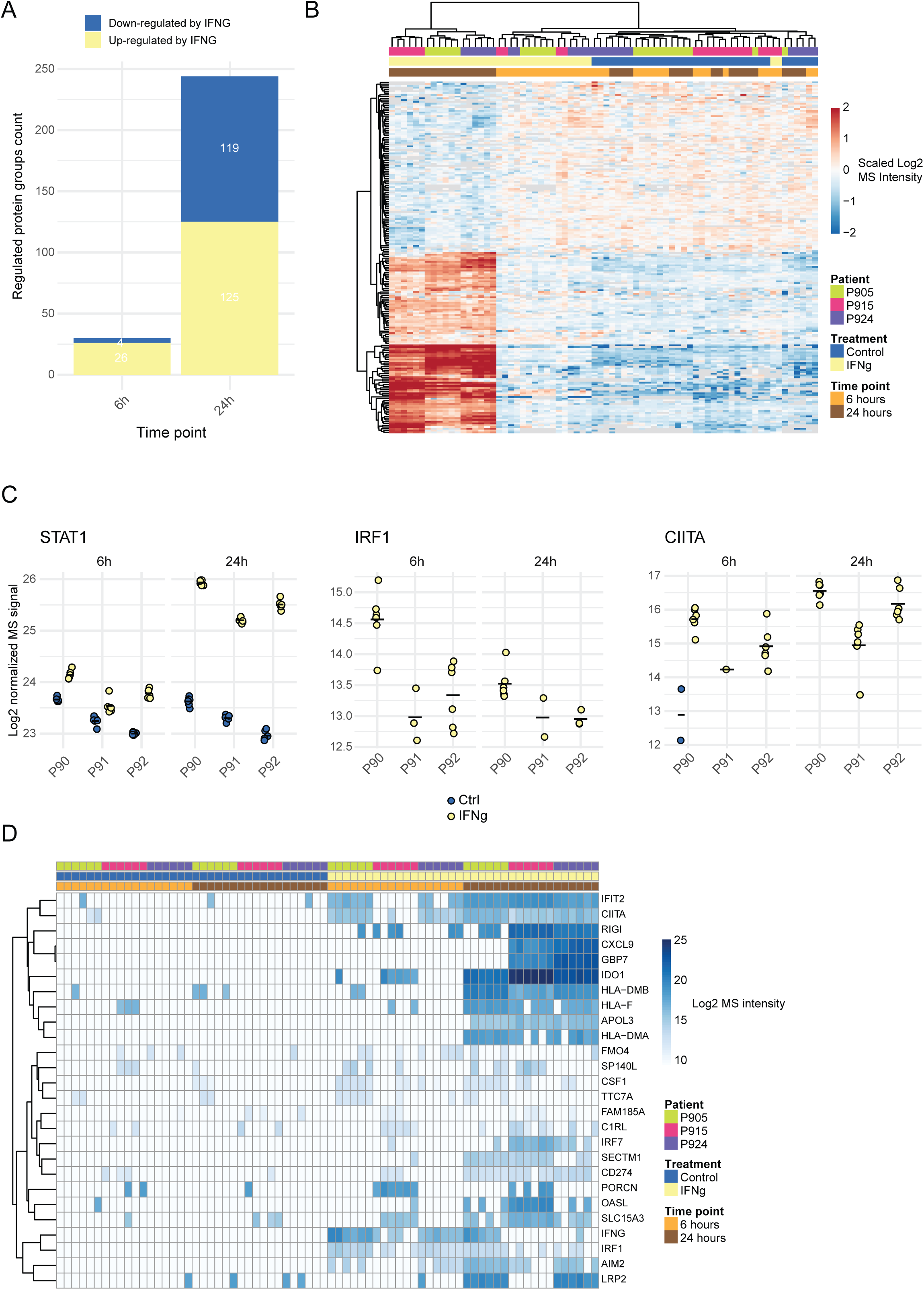
Significantly regulated proteins by IFN-γ. **A**. Number of up- and down-regulated protein groups by IFN-γ at each time point. **B**. Heatmap of scaled log2 MS intensities for protein groups significantly regulated by IFN-γ in at least one time point and with more than 50% of valid values across the entire dataset (n=201 protein groups). Scaling was performed within each protein group and patient. Both rows and columns were clustered with euclidean distance. **C**. Log2 MS intensities of selected protein groups. The black dash represents the mean. **D**. Heatmap of log2 MS intensities of the 38 proteins preferentially expressed upon co-culture, defined as having at most 2 out of 36 valid values in the control condition and at least 7 out of 36 in the IFN-γ condition. Missing values were imputed with a low value. Rows were clustered with euclidean distance, while the column order was fixed.

**Figure S9.**
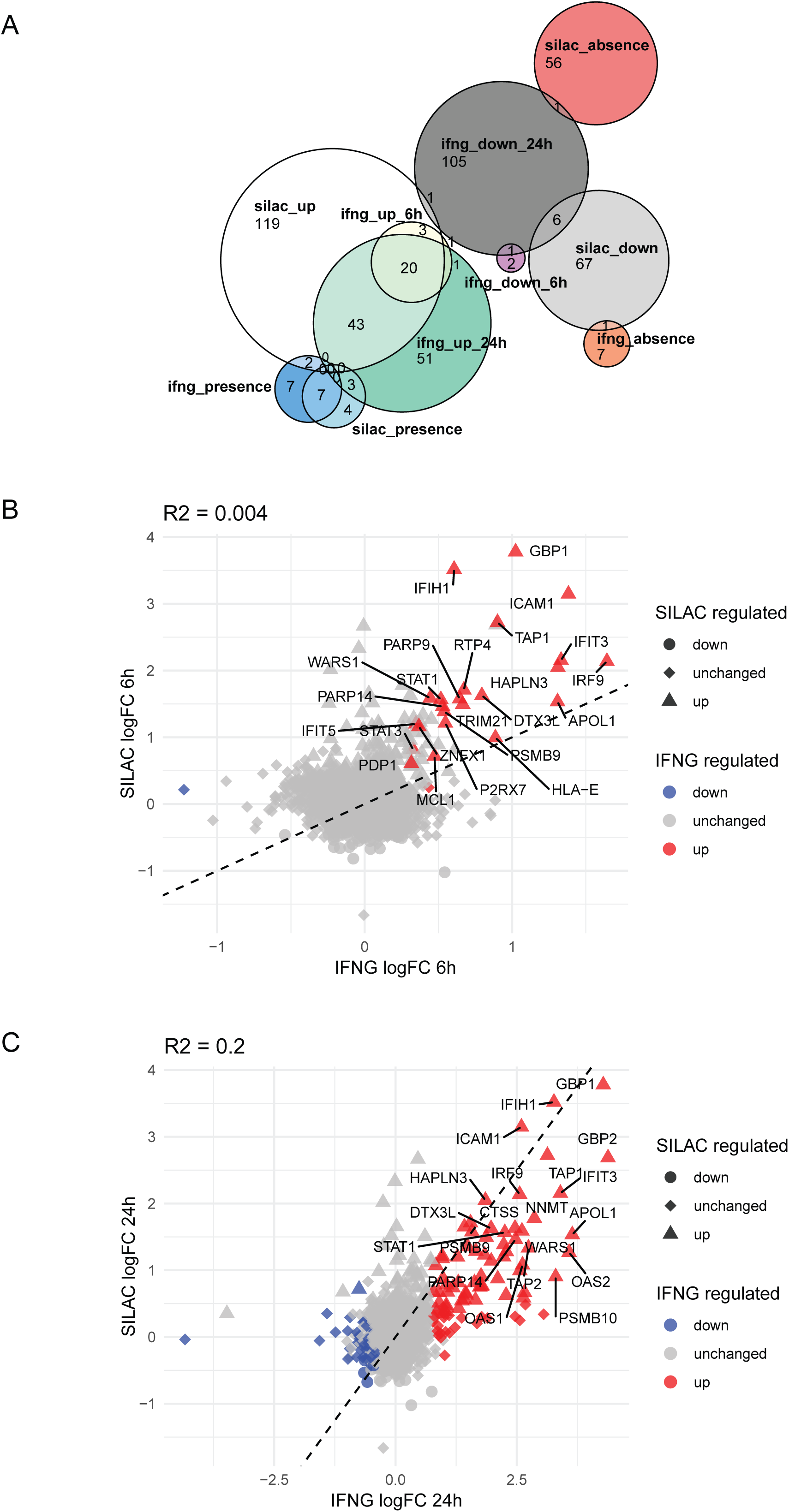
Deconvolution of the IFN-γ-dependent proteome changes upon T cell attack. Related to Figure 3C. **A**. Euler diagram showing the overlap between proteins regulated upon co-culture in the medium-heavy channel and by IFN-γ. **B-C**. Scatter plots comparing the protein fold-change upon 6 hours of co-culture in the medium-heavy channel and upon 2 (B) and 6 hours (C) of IFN-γ. Labeled proteins are shared between the two datasets.

**Figure S10.**
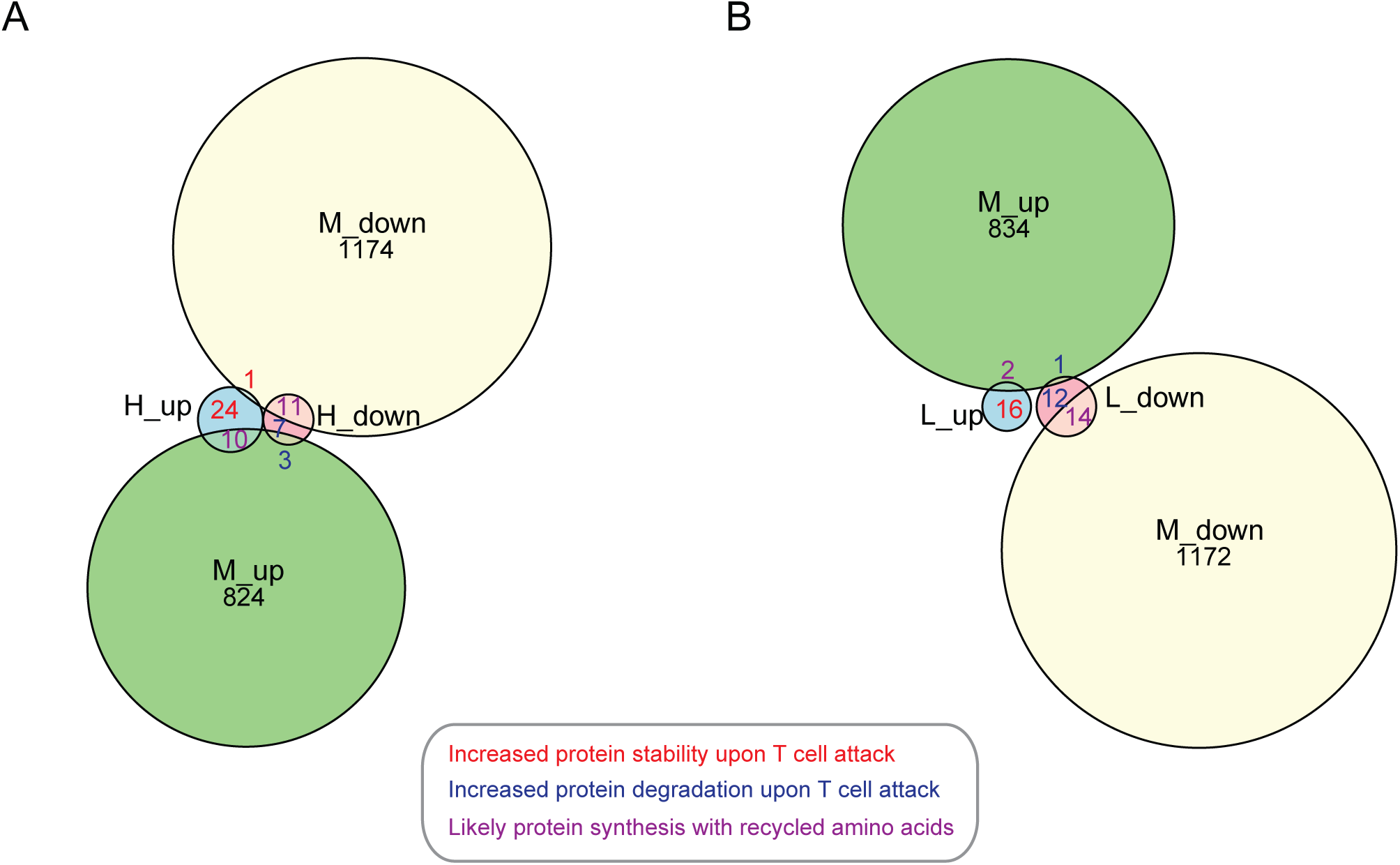
Cell type-resolved analysis of protein turnover upon T cell attack. Related to Figure 4. **A-B**. Euler diagrams showing the overlap between significantly regulated protein groups upon co-culture in the heavy (A) or light channel (A) and protein groups regulated (p <= 0.05) in the medium channel.

**Figure S11.**
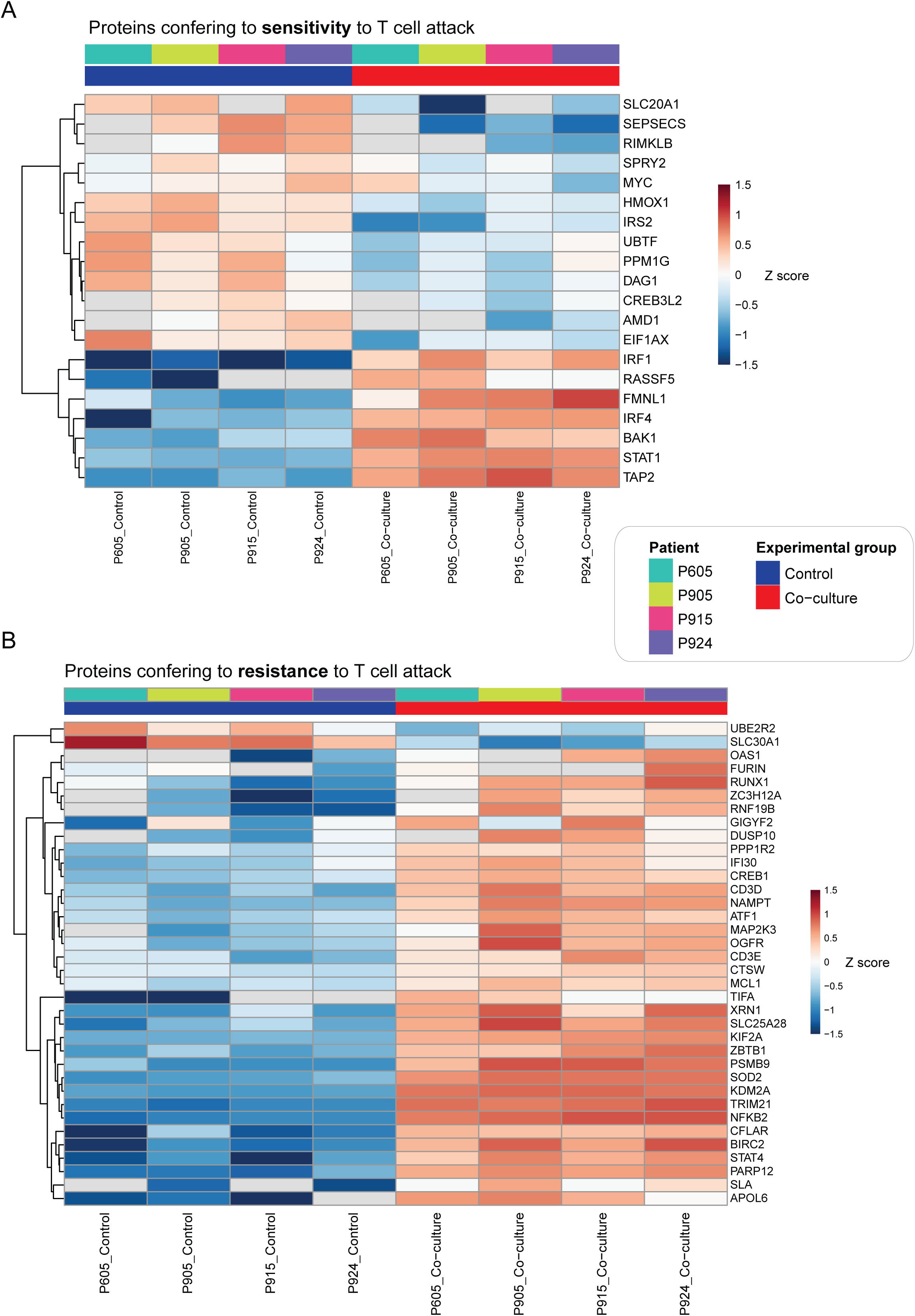
Linking proteomic responses to functional outcomes of T cell attack. **A-B**. Heatmaps of Z-scored log2 MS intensities for proteins conferring sensitivity (A) or resistance (B) to T cell attack. Proteins displayed in A either reduce melanoma cell survival under T cell pressure (enriched in pooled CRISPR-KO screens) and are also significantly upregulated upon attack, or else improve melanoma cell survival under T cell attack (depleted in pooled CRISPR-KO screens) and are also significantly downregulated upon attack. Proteins displayed in B either improve melanoma cell survival under T cell pressure (depleted in pooled CRISPR-KO screens) and are also significantly upregulated upon attack, or else reduce melanoma cell survival under T cell attack (enriched in pooled CRISPR-KO screens) and are also significantly downregulated upon attack. Rows were clustered with euclidean distance, while the column order was fixed.

**Figure S12.**
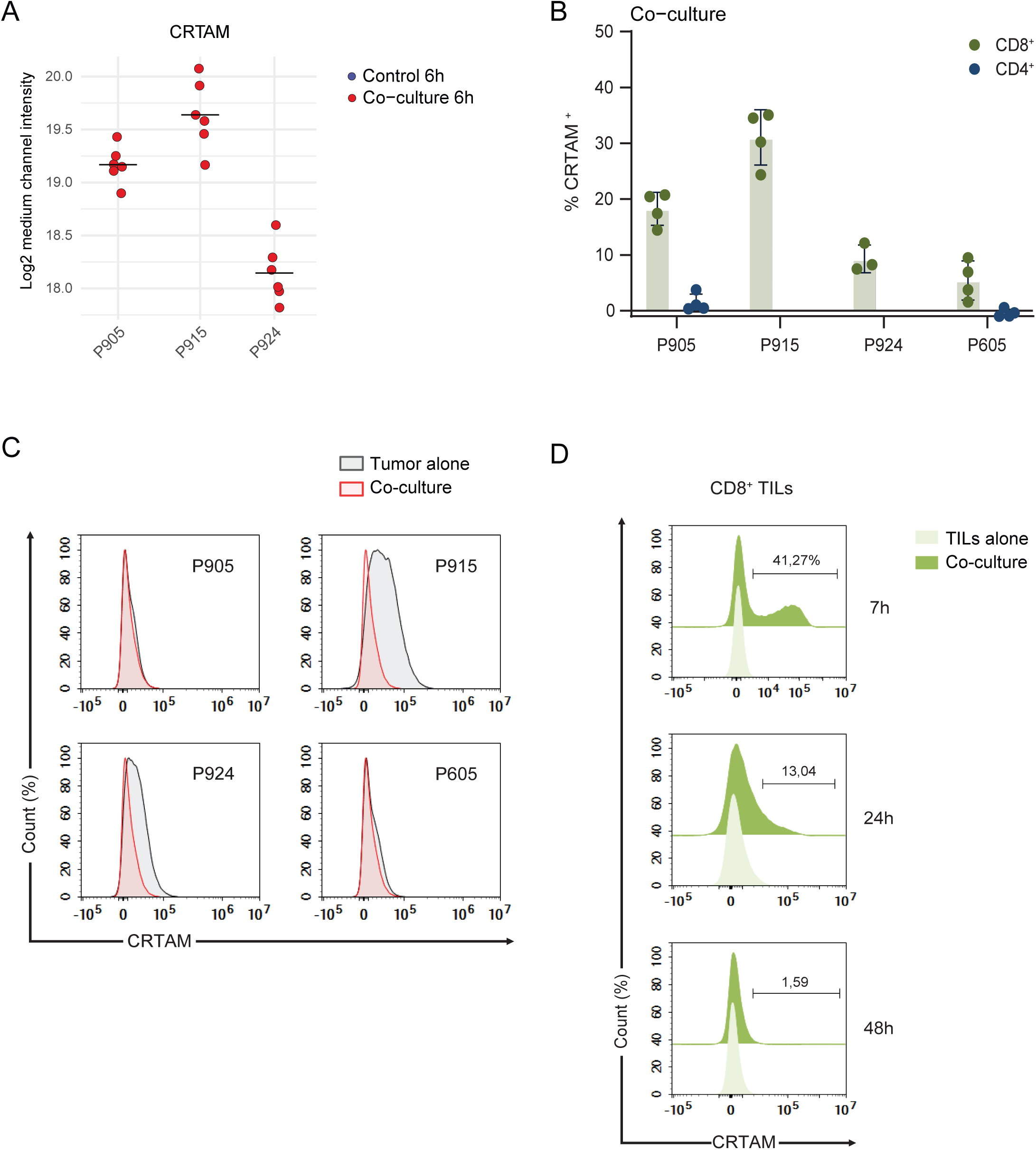
CRTAM expression dynamics in TILs and melanoma cells during co-culture. Related to Figure 5A-B. **A**. Log2 MS intensities of CRTAM in the medium-heavy channel. The black dash represents the mean. **B**. Percentage of CRTAM⁺ TILs in CD4⁺ and CD8⁺ populations across all patients after 6 hours of co-culture with autologous melanoma cells at an E:T ratio of 1:1. No CD4⁺ TILs are shown for P915 and P924, as these REP TIL pools contain negligible CD4⁺ TILs (see Figure S2D). Data are presented as mean ± SD (n = 4 biological replicates, each with three technical replicates). **C**. Histograms showing absence of CRTAM expression on tumor cells alone or after 6 hours of co-culture with autologous TILs at an E:T ratio of 1:1. **D.** Time-course flow cytometry analysis of CRTAM expression in patient 915. CRTAM is strongly upregulated after 7 hours of co-culture with autologous TILs at an E:T ratio of 1:1, followed by a gradual decrease, with expression disappearing by 48 hours post co-culture.

**Figure S13.**
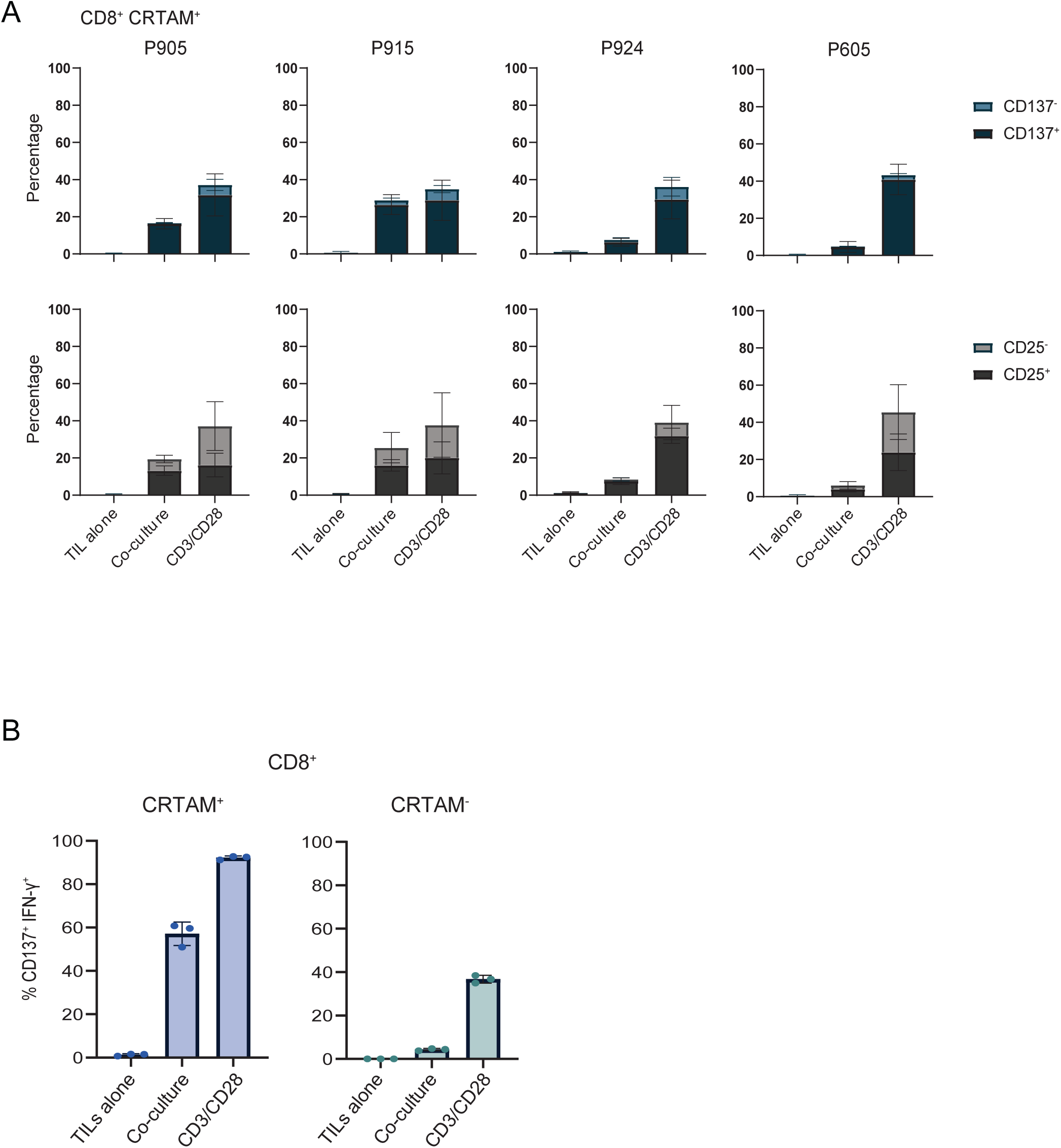
CRTAM⁺ CD8⁺ TILs exhibit enhanced activation and IFN-γ production. Related to Figure 5C-D. **A**. Flow cytometric analysis showing surface expression of CRTAM⁺CD137⁺ versus CRTAM⁺CD137⁻ (top) and CRTAM⁺CD25⁺ versus CRTAM⁺CD25⁻ (bottom) on CD8⁺ TILs across all patients. Cells were analyzed as TILs alone, post 6 hours of CD3/CD28 bead stimulation and co-culture with autologous melanoma cells at an E:T ratio of 1:1. Data are presented as mean ± SD (n = 2 or 4 biological replicates, each with three technical replicates). Complementary to Figure 5C. **B**. Percentage of CD137⁺IFN-γ⁺ co-expression in CRTAM⁺ versus CRTAM⁻ CD8⁺ TILs from patient 915 after 8 hours of co-culture with autologous melanoma cells at an E:T ratio of 1:1 or with CD3/CD28 beads. Gates were set using unstimulated TILs. Data are presented as mean ± SD (n = 3 technical replicates). Complementary to Figure 5D.

**Figure S14.**
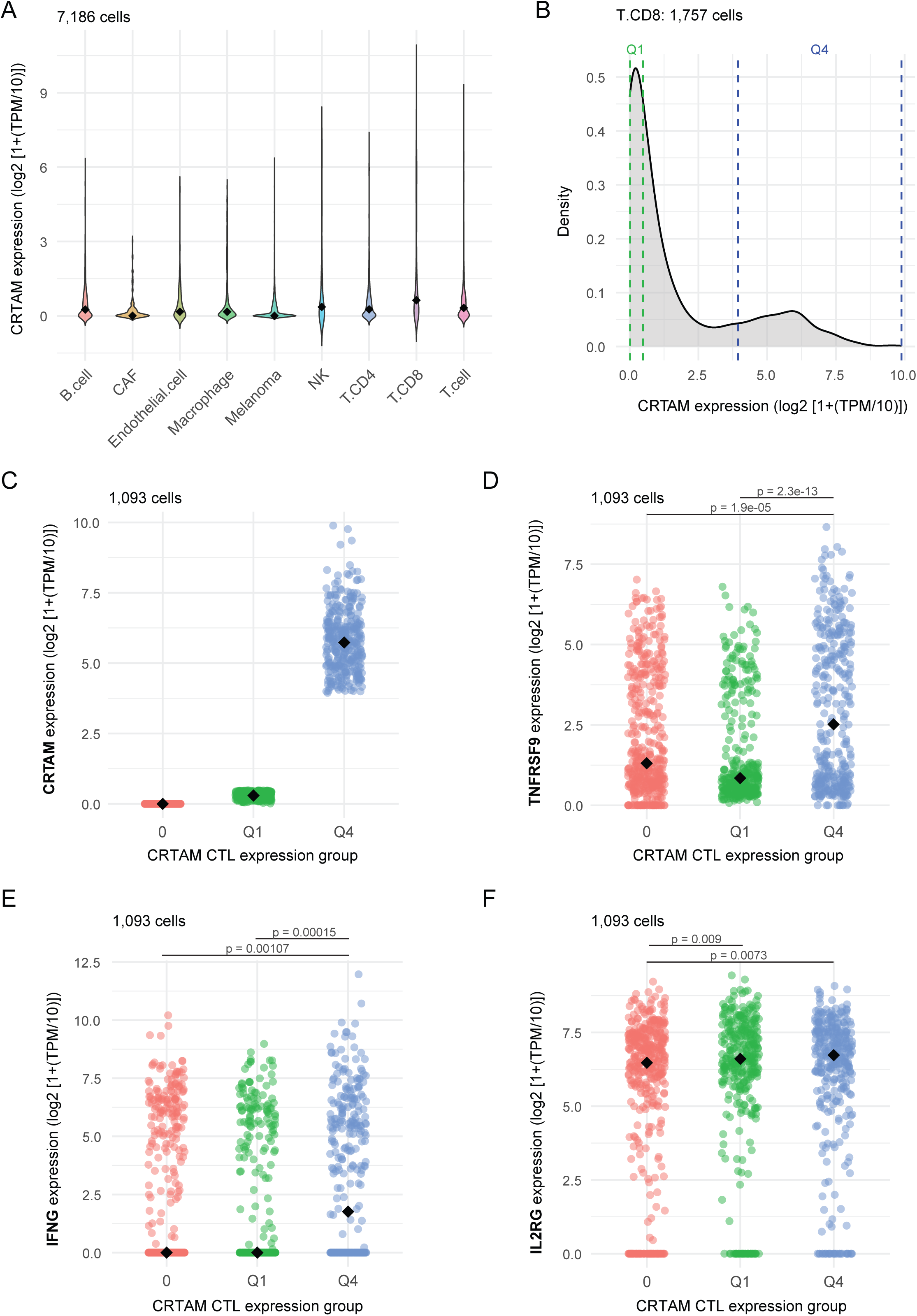
Expression of CRTAM and associated genes in a melanoma single-cell dataset. **A**. Violin plot showing CRTAM gene expression at single-cell level across normal and malignant cell types in a melanoma scRNA-seq dataset. **B**. Histogram showing the CRTAM expression distribution at single cell level across CD8⁺ T cells. The first and fourth quartiles are coloured in green and blue, respectively. **C-F**. Single-cell expression of selected genes in CRTAM-defined groups: no expression (0), first quartile (Q1) and fourth quartile (Q4). p values were calculated by pairwise Wilcoxon Rank Sum Test and corrected with the Benjamini-Hochberg procedure.

**Figure S15.**
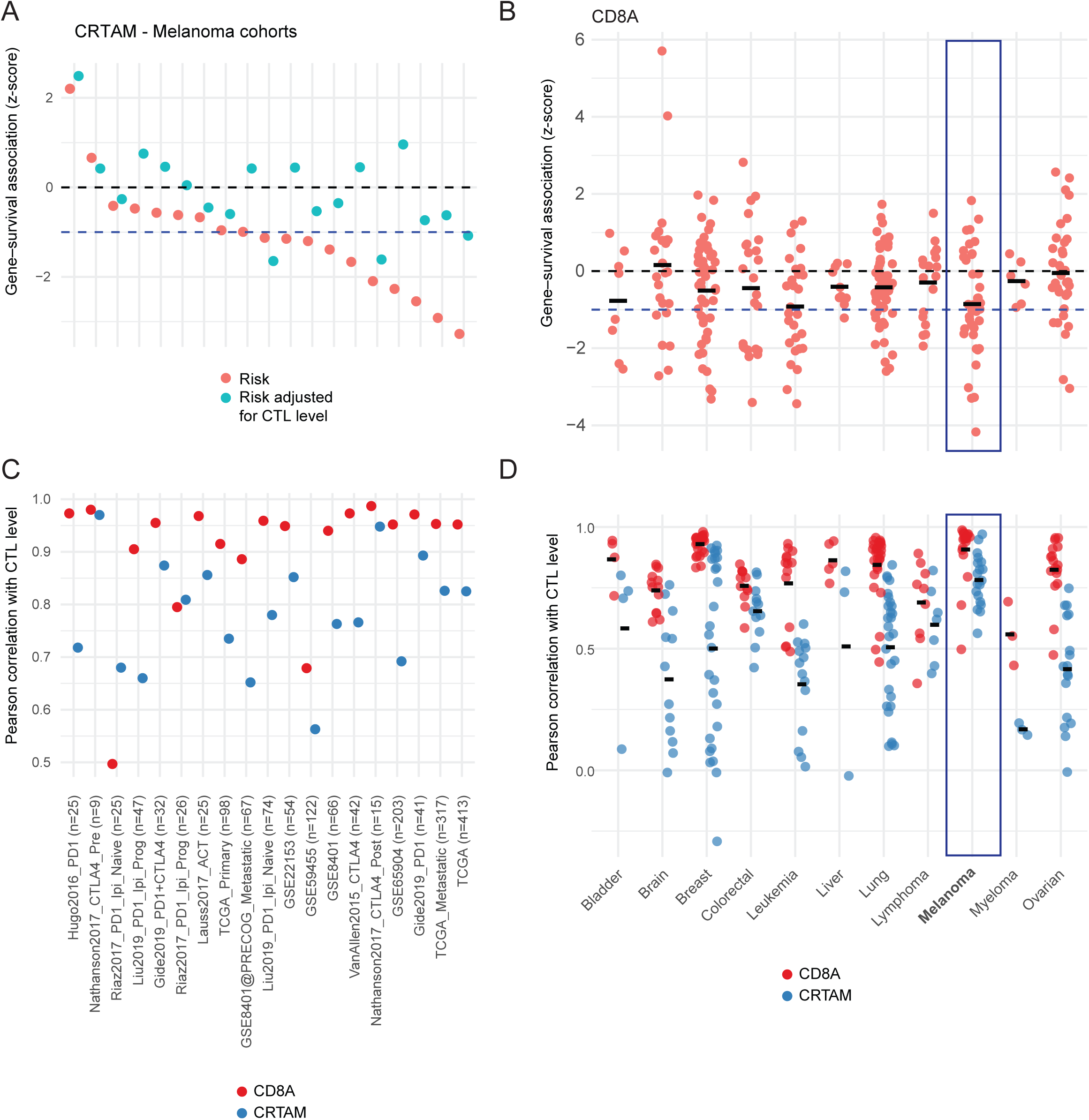
CRTAM analysis in the Tumor Immune Dysfunction and Exclusion (TIDE) database. Related to Figure 5F. **A**. Z-score of the effect of CRTAM gene expression on overall survival across multiple melanoma cohorts in a CoxPH model, before and after adjusting for CTL infiltration. A negative z-score means that higher CRTAM expression is associated with lower death risk. **B**. Z-score of the effect of CD8A gene expression on overall survival across multiple cancer types in a CoxPH model, before and after adjusting for CTL infiltration. Melanoma cohorts are circled in blue. **C-D**. Pearson correlation between CRTAM gene expression and CTL infiltration across multiple melanoma cohorts (C) and cancer types (D). Melanoma cohorts in D are circled in blue.

**Figure S16.**
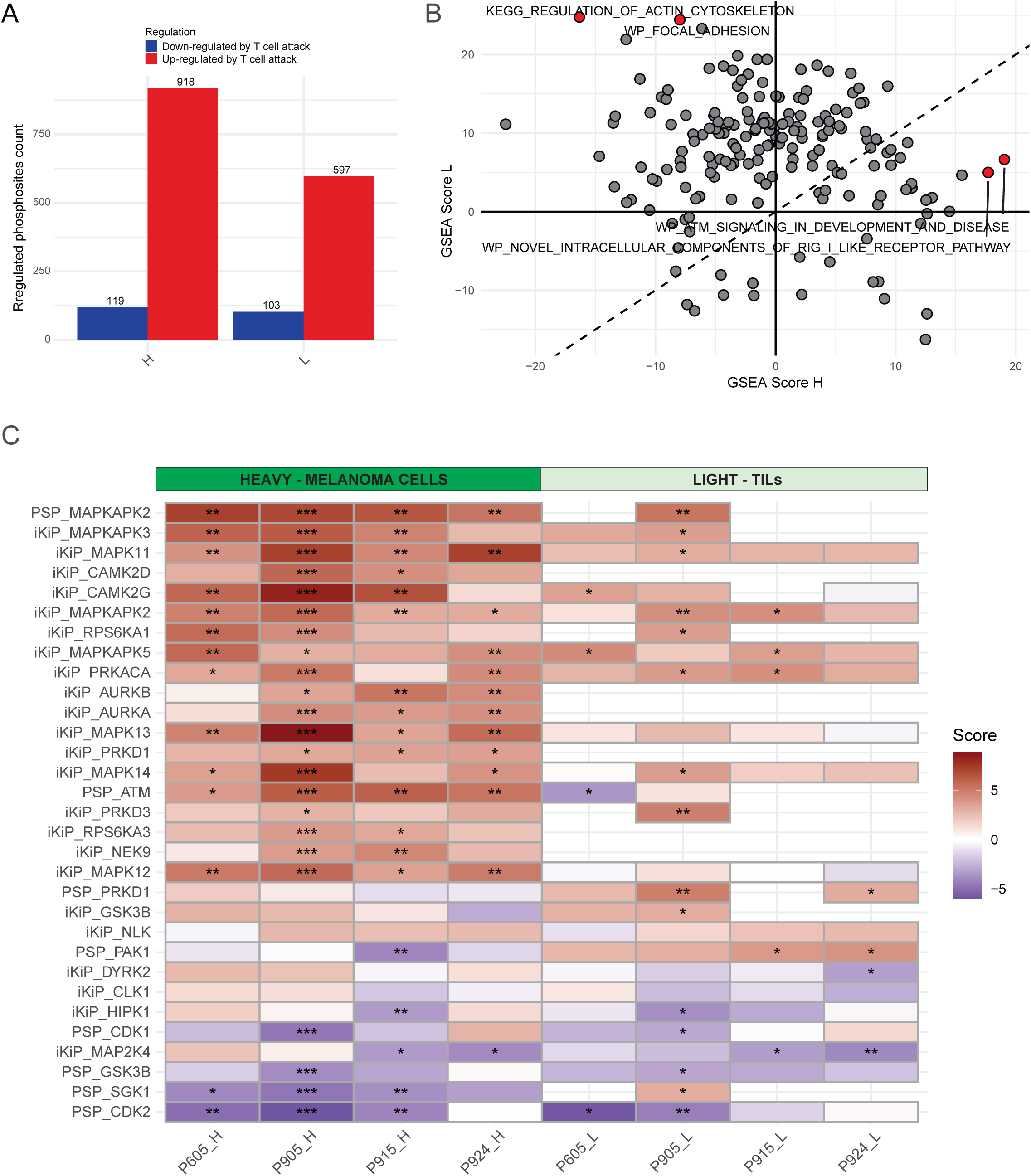
Phosphosites, phosphorylation driven-pathways and kinases regulated upon T cell attack. Related to Figures 6-7. **A**. Number of up- and down-regulated phosphosites per contrast and channel. **B**. Scatter plot of gene-centric-redundant ssGSEA scores in the heavy and light channels. The score is calculated by summing the individual patients’ scores. Only pathways with adjusted p in at least one patient in one channel are shown. **C**. Heatmaps of PTM-SEA kinase scores. Only kinases expressed on the proteome level are shown. The 20 kinases with the highest sum of -log10(adjusted p) across the four patients per channel were selected for this plot, for a total of 31 unique kinases. Significance is represented by asterisks: * = adjusted p <= 0.05; ** = adjusted p <= 0.01; *** = adjusted p <= 0.005.

**Figure S17.**
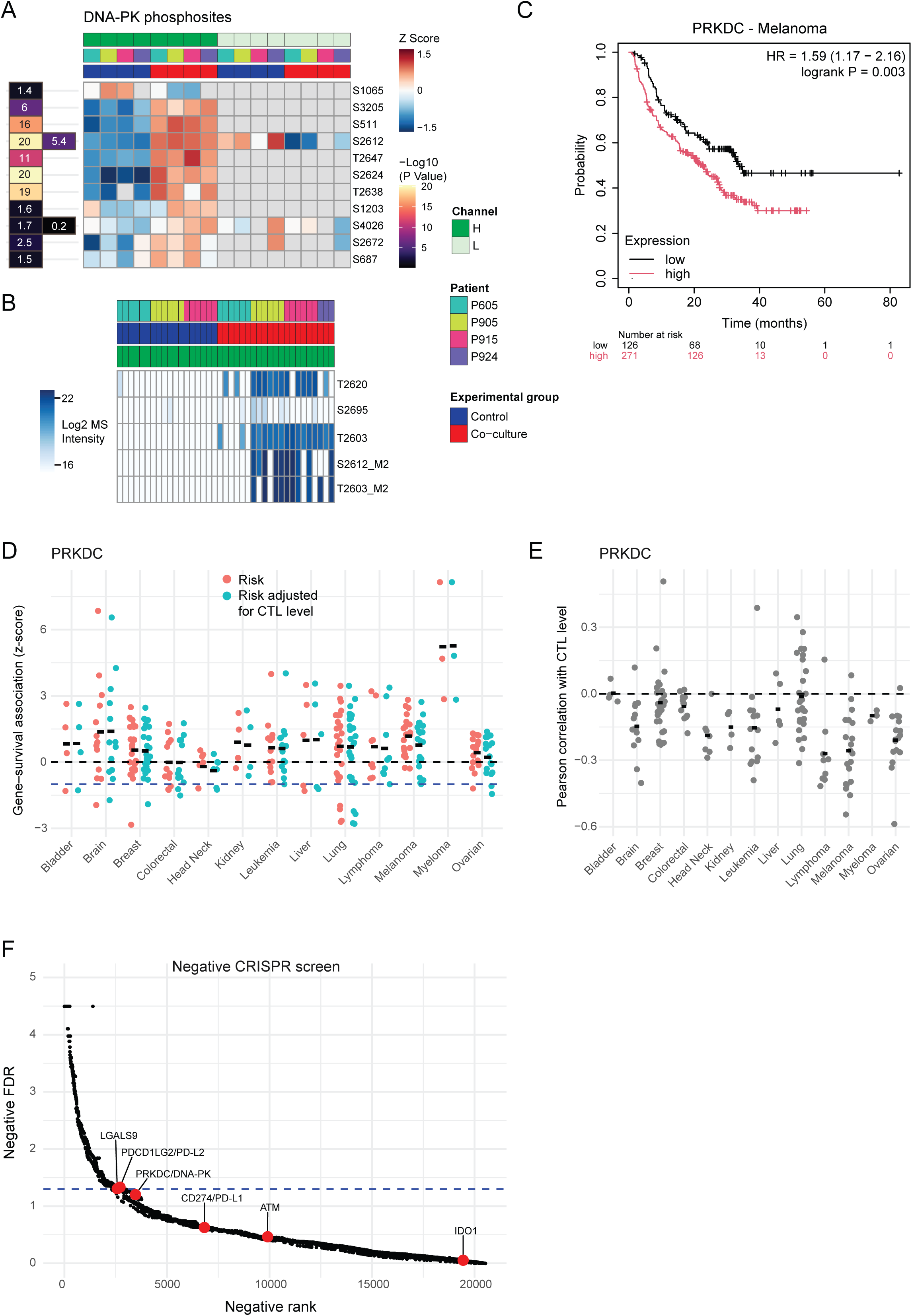
DNA-PK: a potential immune resistance kinase in melanoma. **A**. Heatmaps of Z-scored log2 MS intensities (right) and -log10(p value) (left) for PRKDC phosphosites with a p value <= 0.05. **B**. Heatmap of log2 MS intensities of the 5 PRKDC phosphosites preferentially expressed upon co-culture, defined as having at most 2 out of 48 values in the control condition and at least 10 out of 48 in the co-culture condition. Missing values were imputed with a low value. **C**. KM curve showing the association between PRKDC gene expression and survival in a cohort of 423 melanoma patients undergoing treatment with immune checkpoint inhibitors. **D**. Z-score of the effect of PRKDC gene expression on overall survival across multiple cancer types in a CoxPH model, before and after adjusting for CTL infiltration. A positive z-score means that higher PRKDC expression is associated with higher death risk. Melanoma cohorts are circled in blue. **E**. Pearson correlation between PRKDC gene expression and CTL infiltration across multiple cancer types. Melanoma cohorts are circled in blue. **F**. Negative FDR as a function of negative rank from the CRISPR-KO data of Zhang et al. Some of the most well-known immune checkpoint molecules are labeled in red.

